# Neonatal systemic gene therapy restores cardiorespiratory function in a rat model of Pompe disease

**DOI:** 10.1101/2024.12.10.627800

**Authors:** David D. Fuller, Sabhya Rana, Prajwal Thakre, Ethan Benevides, Megan Pope, Adrian G. Todd, Victoria N. Jensen, Lauren Vaught, Denise Cloutier, Roberto A. Ribas, Reece C. Larson, Matthew S. Gentry, Ramon C. Sun, Vijay Chandran, Manuela Corti, Darin J. Falk, Barry J. Byrne

## Abstract

Absence of functional acid-α-glucosidase (GAA) leads to early-onset Pompe disease with cardiorespiratory and neuromuscular failure. A novel Pompe rat model (*Gaa^-/-^*) was used to test the hypothesis that neonatal gene therapy with adeno-associated virus serotype 9 (AAV9) restores cardiorespiratory neuromuscular function across the lifespan. Temporal vein administration of AAV9-DES-GAA or sham (saline) injection was done on post-natal day 1; rats were studied at 6-12 months old. Whole-body plethysmography showed that reduced inspiratory tidal volumes in *Gaa^-/-^*rats were corrected by AAV-GAA treatment. Matrix-assisted laser desorption/ionization mass spectrometry imaging (MALDI) revealed that AAV-GAA treatment normalized diaphragm muscle glycogen as well as glycans. Neurophysiological recordings of phrenic nerve output and immunohistochemical evaluation of the cervical spinal cord indicated a neurologic benefit of AAV-GAA treatment. *In vivo* magnetic resonance imaging demonstrated that impaired cardiac volumes in *Gaa^-/-^* rats were corrected by AAV-GAA treatment. Biochemical assays showed that AAV treatment increased GAA activity in the heart, diaphragm, quadriceps and spinal cord. We conclude that neonatal AAV9-DES-GAA therapy drives sustained, functional GAA expression and improved cardiorespiratory function in the *Gaa^-/-^* rat model of Pompe disease.

## INTRODUCTION

Pompe disease results from mutations in the gene encoding acid-α-glucosidase (GAA), an enzyme essential for the degradation of lysosomal glycogen. A recent evaluation of >11M newborns screened for Pompe disease indicates that prevalence is approximately 1 per 19,000 births.^1^ Infantile-onset Pompe disease (IOPD) patients lack functional GAA protein and experience cardiorespiratory failure if untreated.^2^ Deficiency in GAA results in widespread glycogen accumulation and disruption of cellular architecture and function in cardiac, skeletal, and smooth muscle, as well as neurons, particularly motor neurons.^3,4^

The standard treatment for Pompe disease is enzyme replacement therapy (ERT), which requires biweekly intravenous infusion of recombinant human GAA protein. ERT has high medical costs, and patients with little or no residual GAA expression often respond poorly. Furthermore, ERT does not effectively target the central nervous system (CNS) pathology^5–8^ , a critical consideration given the growing evidence of CNS involvement in Pompe disease ^5,9^, including the potential contribution to respiratory failure.^10–13^ Consequently, ERT has limited success in preventive respiratory failure.^4^ A clinical trial targeting the diaphragm muscle^14,15^ as well as data from mouse models^16–18^ indicate that gene therapy using adeno-associated virus (AAV) is a viable approach to treat Pompe disease. A particular advantage of AAV-GAA therapy is the ability to effectively target skeletal and cardiac muscle as well as the CNS.^19^

In this study we assessed the effectiveness of an AAV gene therapy to treat Pompe pathology in a new *Gaa*- null rat model of Pompe disease, which exhibits neuromuscular glycogen accumulation and cardiorespiratory dysfunction. We tested the hypothesis that initiating gene therapy in neonates would lead to permanent GAA expression in the CNS, cardiac, and skeletal muscles, ultimately improving cardiorespiratory function. We utilized an AA serotype 9 (AAV9) vector, encoding human GAA (hGAA) driven by a desmin (DES) promoter. AAV9 was selected as previous work has demonstrated its ability to effectively drive cardiac and neuronal gene expression.^16,20^ A comprehensive battery of outcomes measures included cardiac imaging, plethysmographic and neurophysiologic assessment of breathing, histology, and matrix-assisted laser desorption/ionization mass spectrometry imaging (MALDI-MSI^21^) of tissues. The results demonstrate that neonatal AAV9-DES-GAA therapy successfully drives sustained, functional GAA expression and prevents cardiorespiratory decline in the *Gaa^-/-^* rat model of Pompe disease.

## RESULTS

Breathing patterns (*e.g*., **Figure 1A**) and body weight were measured repeatedly from six to twelve months of age. These studies showed that AAV-GAA treatment had a profound effect on body weight as the rats aged (treatment, P<0.001; **Figure 1B**). Thus, weight was similar between Sprague-Dawley (S-D) and AAV- GAA treated *Gaa^-/-^* rats but was substantially reduced in the phosphate buffered saline treated *Gaa^-/-^*rats. Whole body plethysmography measurements showed that inspiratory tidal volume (VT, ml/br) was greater in the AAV-GAA treated as compared to the saline-treated *Gaa^-/-^* rats (treatment, P<0.001, **Figure 1C**). A significant group effect was also observed for minute ventilation (V̇E, ml/min; P=0.003, **Figure 1D**). Respiratory rate (breaths per minute) showed a strong trend to be different across groups, most notably versus the saline-treated *Gaa^-/-^* rats breathing at higher rates at 6- and 9-mo compared to the gene therapy group (treatment, P=0.057, **Figure 1E**).

**Figure 1.**
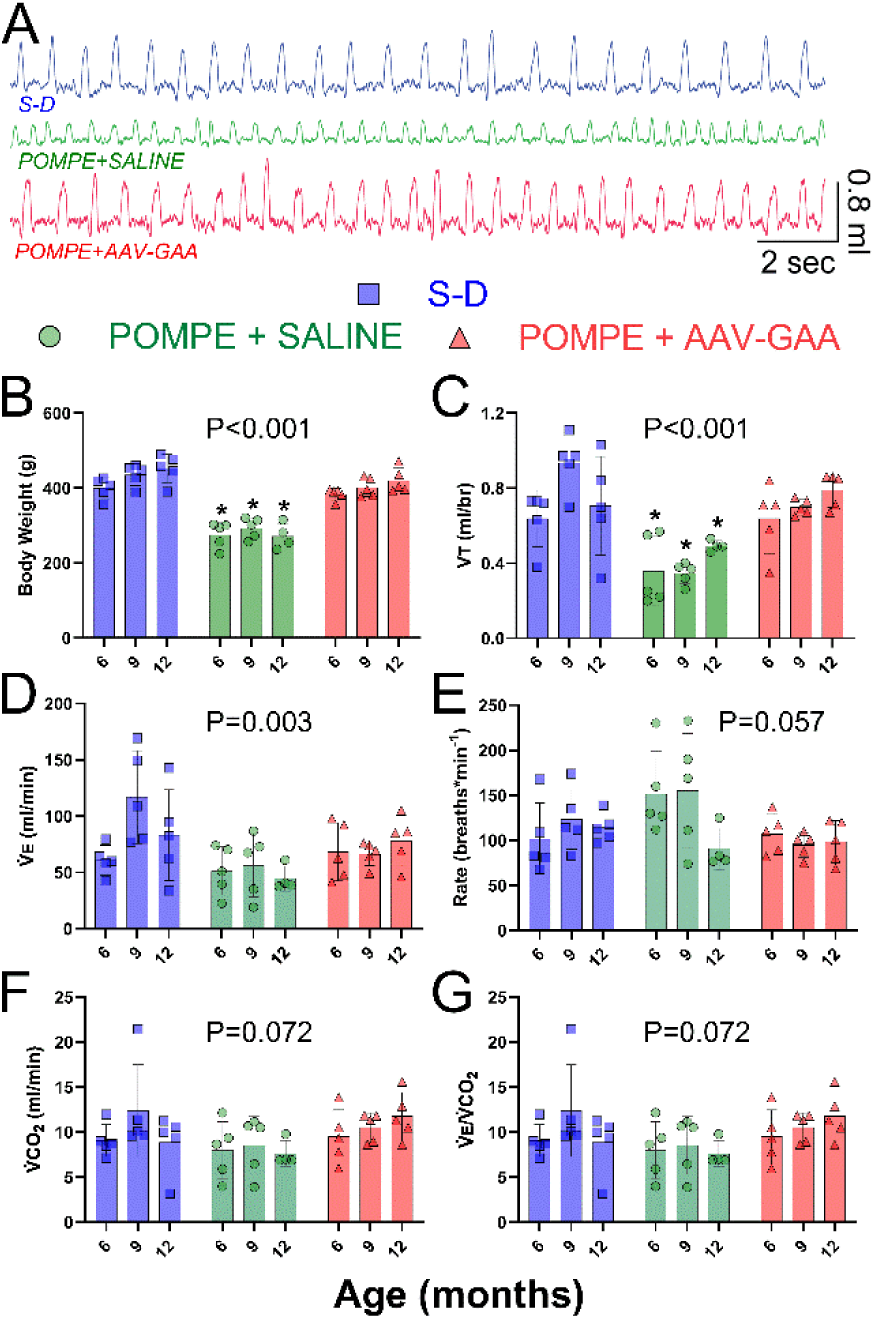
AAV-GAA treatment normalizes body weight and breathing. Data obtained during baseline room air breathing *i.e*., “eupnea”. **A:** Example of breathing patterns measured using whole body plethysmography. **B:** Body weight is normalized after AAV treatment. **C:** Tidal volume (VT, ml/br) is restored after AAV treatment. **D:** Minute ventilation (V̇E, ml/min) is restored after AAV treatment. Respiratory rate (**E**), metabolic rate (as estimated via V̇CO_2_; **F**) and the ratio of V̇E to V̇CO_2_ (**G**) all showed a strong trend to be impacted by AAV treatment. Statistical test: 2-way RM ANOVA. The treatment effect P-value is reported on each plot. *, p<0.05 vs. Pompe + AAV-GAA. S-D: Sprague-Dawley

We also observed that metabolic rate, as estimated by CO_2_ production measured in the plethysmography chamber (*i.e.*, V̇CO_2_), tended to be enhanced in *Gaa^-/-^* rats that received AAV-GAA treatment (treatment, P=0.072, **Figure 1F**). The ratio of minute ventilation to metabolic rate (V̇E /V̇CO_2_) was consistent with hypoventilation in saline-treated *Gaa^-/-^* rats. **Figure 1G** shows that the V̇E /V̇CO_2_ was reduced in *Gaa^-/-^* rats compared to *Gaa^-/-^*rats following AAV-GAA treatment or Sprague-Dawley rats (treatment, P=0.072). Breathing was also studied during an acute respiratory challenge achieved by brief exposure to a 10% O_2_, 7% CO_2_ gas mixture. These data are shown in **Figure S1** and indicate that the ability to increase VT and V̇E during a breathing challenge was restored by AAV-GAA treatment in *Gaa^-/-^* rats (treatment, P=0.003).

Continuing the focus on the respiratory system, the spatial metabolomic profile of the diaphragm muscle was analyzed *ex vivo* at age 12 months using MALDI mass spectrometry imaging (MSI); example diaphragm tissue images are shown in **Figure 2A**. First, MALDI-MSI revealed a major reduction in diaphragm glycogen quantified by glycogen chain length abundance (cleaved with isoamylase, see methods) in *Gaa^-/-^*rats that received the neonatal AAV-GAA treatment (**Figure 2B**). Since glycogen is directly channeled to central carbon metabolism of bioenergetics, lipid biosynthesis, and complex carbohydrate metabolism such as N-linked glycan^22–25^, we examined metabolome, lipidome, as well as glycome using MALDI^22–26^. We performed hierarchical clustering based on similarity of individual biological replicates. Excitingly, treatment of AAV-GAA in the *Gaa^-/-^* animals provided lifelong normalization of the diaphragm glycome as shown by the unsupervised clustering analysis, i.e. inability to separate WT or *Gaa^-/-^* treated with AAV animals by hierarchical clustering (**Figure 2C)**. A similar clustering heatmap analysis is performed for metabolomics datasets, and in agreement with glycome analysis, metabolomics and lipidomics analyses also showed the strong impact of AAV-GAA therapy on the diaphragm to return to WT metabolic profiles. (**Figure 2D**). Representative molecules are displayed to show strong rescue phenotype after AAV treatment in the *Gaa^-/-^*animals (**Figure 2E-J)**. For example, diaphragm levels of glucose, 3-phosphoglyceric acid (3PG), glycerophosphorylethanolamine, docosahexaenoic acid, and arachidonic acid were all normalized in the AAV-GAA treated *Gaa^-/-^* rats.

**Figure 2.**
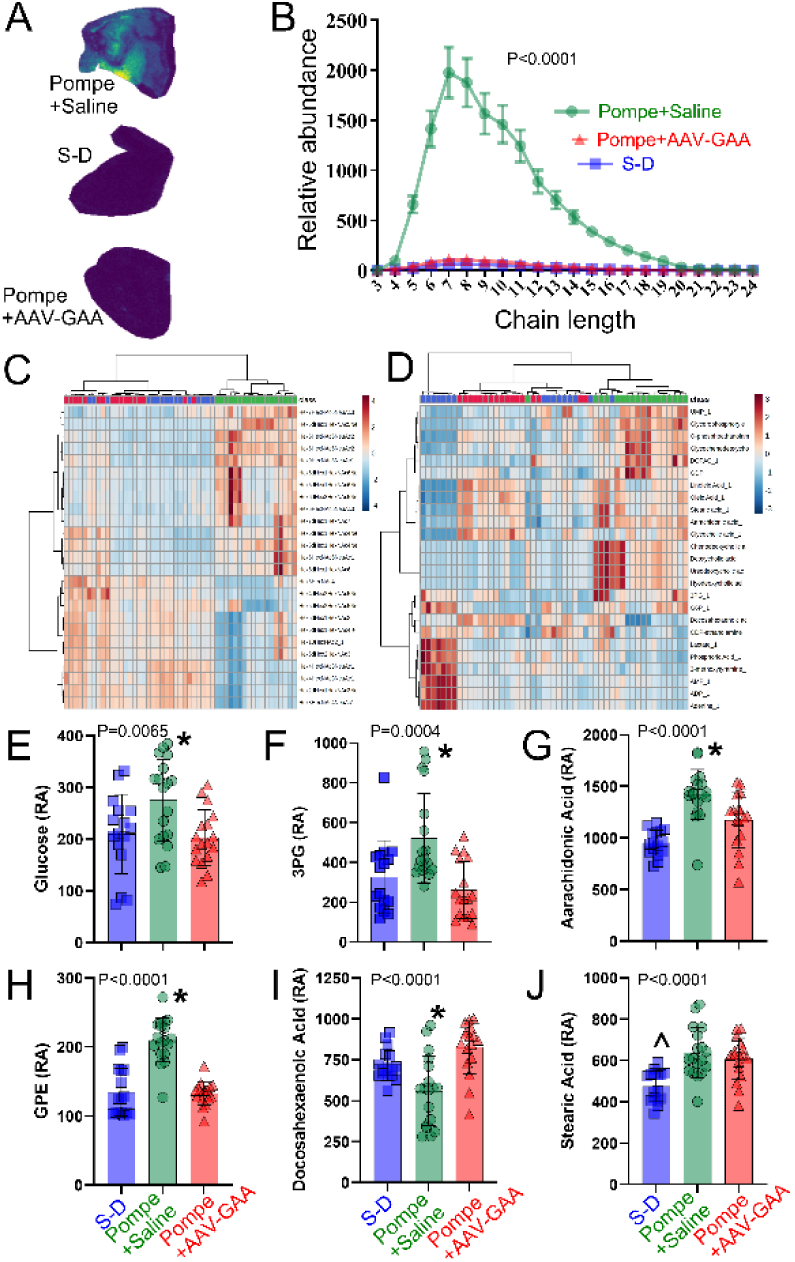
Spatial metabolomic profile of the diaphragm muscle at age 12 months indicates normalization of glycogen after AAV-GAA. A: Examples of diaphragm tissue evaluated using MALDI. The heat map shows the gradient of glycogen in diaphragm (represented by chain length +7, 1175m/z). **B:** Normalization of diaphragm glycogen after neonatal AAV-GAA treatment. **C-D**: Unsupervised clustering heatmap analysis for the glycome (**C**) and metabolome/lipidome (**D**). the treatment group is indicated by the top row, and the relative expression of each molecule is indicated by the color on the heat map. Relative abundance (RA) plots show glucose (**E**), 3-Phosphoglyceric acid (3PG) (**F**), arachidonic acid (**G**), glycerophosphorylethanolamine (GPE) (**H**), docosahexaenoic acid (**I**), and stearic acid (**J**). Statistical tests: **B:** 2-way RM ANOVA; treatment effect P-value is reported on plot. **E-J:** 1-way ANOVA; treatment effect P-value is reported on each plot. *, p<0.05 vs. Pompe+AAV-GAA; S-D: Sprague-Dawley. Color scheme for treatment groups is the same on all panels.

The diaphragm was histologically evaluated in a separate cohort of rats at six months of age; example photomicrographs are shown in **Figure 3A**. Saline-treated *Gaa^-/-^* rats had a reduction in the size (cross-sectional area, CSA) of Type I (treatment, P=0.002) and Type IIb/x diaphragm myofibers (treatment, P<0.001) as compared to Sprague-Dawley rats (**Figure 3B**). However, myofiber size was normalized in *Gaa^-/-^* rats following the AAV-GAA treatment, with values comparable to that observed in Sprague-Dawley rats. We also observed an impact of AAV-GAA treatment on the overall number of diaphragm Type IIb/x fibers (treatment, P<0.001) as shown in **Figure 3C**.

**Figure 3.**
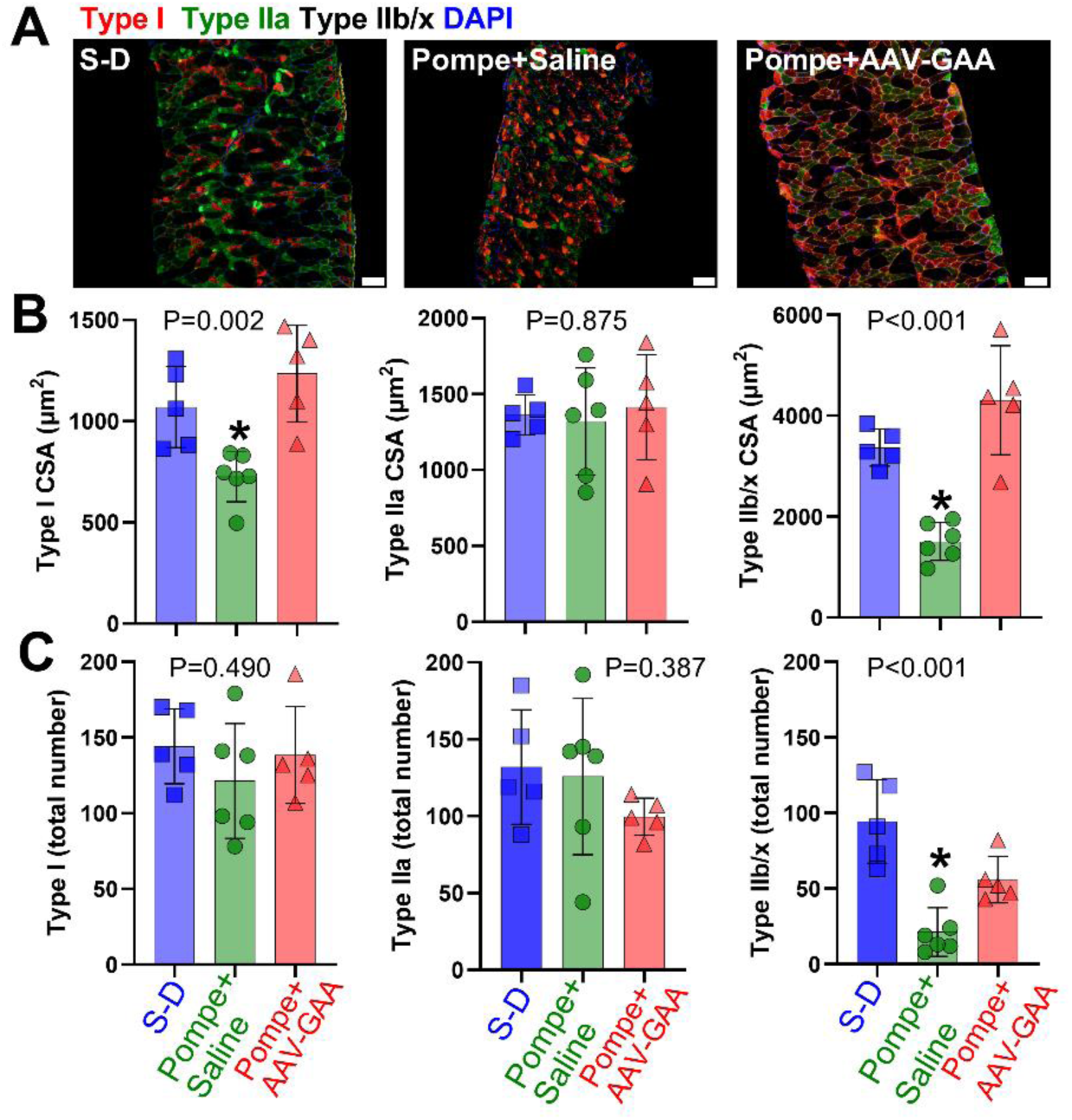
Impact of AAV-GAA treatment on diaphragm myofibers. A: Example photomicrographs from each group. **B:** Type I and IIb/x myofiber size was normalized in Pompe rats following the AAV- GAA treatment, with values comparable to that observed in Sprague-Dawley (S-D) rats. **C:** AAV-GAA treatment increased the overall number of diaphragm Type IIb/x fibers (treatment, P<0.001). Statistical test: 1-way ANOVA; P-value is reported on each plot. *, p<0.05 vs. Pompe+AAV-GAA

Direct recordings from the phrenic nerve (e.g., **Figure 4A**) were performed in anesthetized rats at 5-6 months of age. Neurophysiology experiments were performed to evaluate if the neural drive to the diaphragm was impacted by AAV-GAA treatment in *Gaa^-/-^*rats. Recordings were performed under controlled conditions in which arterial blood gases were monitored and standardized between groups. All baseline recordings were made with the end-tidal CO_2_ at 4 mmHg above the threshold for evoking inspiratory bursting. We observed that in Pompe rats phrenic motor output was unstable if the CO_2_ values were below this value. As shown in **Figure 4B**, *Gaa^-/-^* rats treated with AAV-GAA had evidence of increased neural drive to the diaphragm as reflected by the amplitude of the inspiratory efferent phrenic burst. This observation however did not reach statistical significance (treatment, P=0.060). Like the breathing data collected in awake rats, the respiratory rate (bursts per minute) was similar between *Gaa^-/-^*saline- and AAV-GAA treated rats (treatment, P=0.806, **Figure 4C**). We also evaluated heart rate (beats per min) during these experiments and observed an elevated rate in AAV-GAA *vs*. saline treated *Gaa^-/-^* rats (treatment, P=0.036).

**Figure 4.**
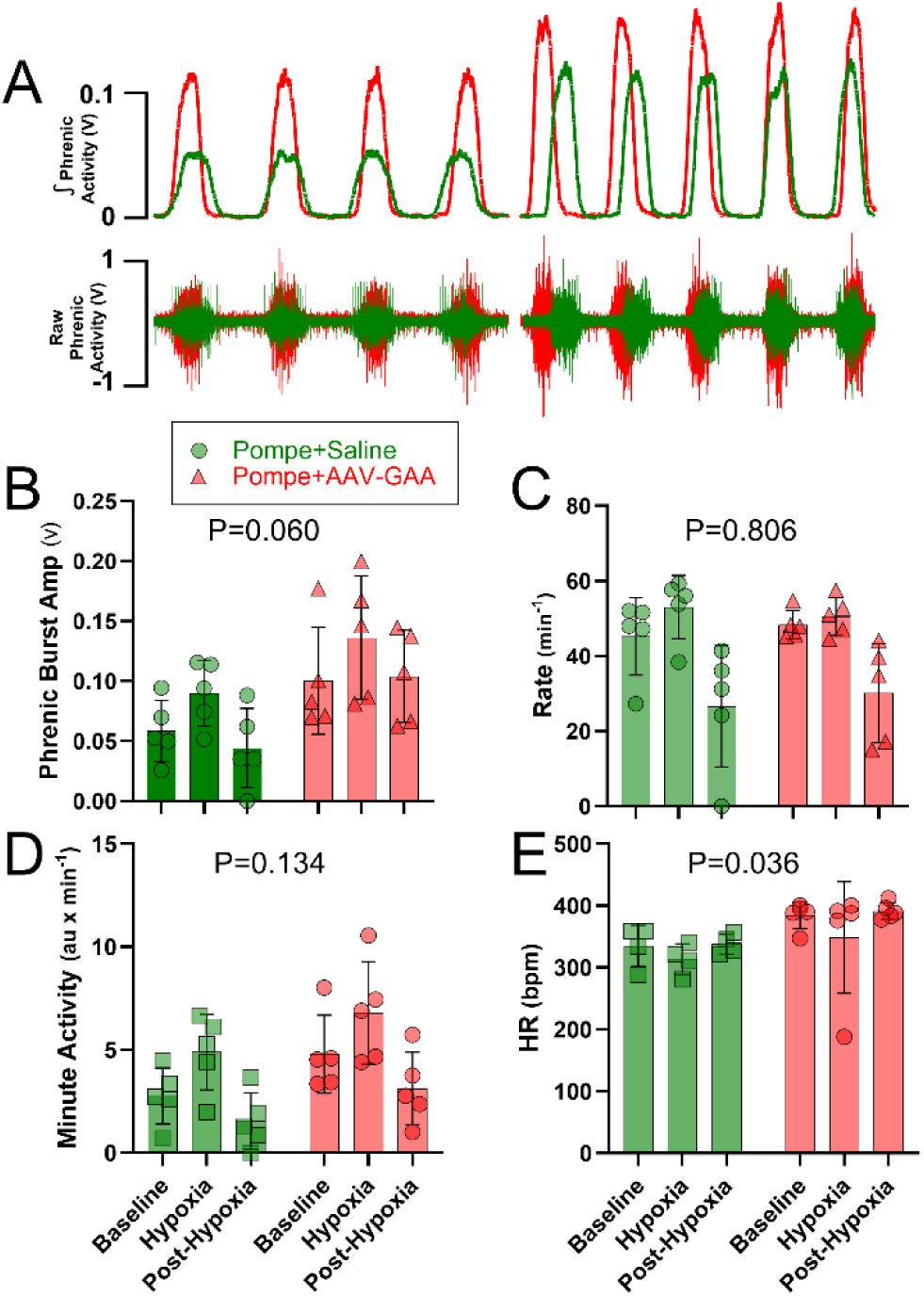
Phrenic nerve recordings. A: Examples of recordings of inspiratory bursting recorded in the phrenic nerve of anesthetized rats during baseline conditions. **B**: Inspiratory burst amplitude (v). Pompe rats treated with AAV-GAA showed a strong trend for increased burst amplitude (P=0.060). **C:** Respiratory rate (bursts per minute) was similar between saline and AAV-GAA treated rats (P=0.806). **D:** Heart rate (beats per min) was greater in AAV-GAA *vs*. saline treated rats (P=0.036). Statistical test: 2-way RM ANOVA. The treatment effect P-value is reported on each plot. S-D: Sprague-Dawley

The suggestion of an impact on efferent phrenic neural output (**Figure 4B**) and the demonstration of larger inspiratory VT after AAV-GAA treatment (**Figure 1B**) led us to histologically examine the spinal cord (n=2 each group), focusing on the region of the phrenic motoneurons that innervate the diaphragm (*e.g.*, mid-cervical ventral horn). A glycogen antibody was used to visually assess the presence and localization of glycogen in the cervical spinal cord (**Figure 5A**). Neurons in the anterior horn of the spinal cord of the *Gaa^-/-^* rat showed the prototypical histopathology that is well established in Pompe disease, including a swollen soma and glycogen accumulation. Further, these neurons displayed large accumulation of glycogen within the cell bodies. On qualitative examination, positive staining for neuronal glycogen was considerably reduced in *Gaa^-/-^* rats that were treated with AAV-GAA (**Figure 5B**). This was evidenced by a reduction in glycogen positive puncta within the soma of motor neurons in the ventral horn of the spinal cord. Tissue sections were also stained for astrocytes (GFAP) and microglia (Iba1). GFAP staining was abundant in the grey matter regions of the anterior horn of the spinal cord. Qualitatively, a higher density of astrocytes was observed near motor neurons in the *Gaa^-/-^* rat. In the AAV-GAA treated rats, a reduction in astrocytes was observed. No apparent differences in microglia were observed between the two groups. **Figure 5C-E** provides higher magnification examples of neurons in the ventral horn of the mid-cervical (C4) spinal cord.

**Figure 5.**
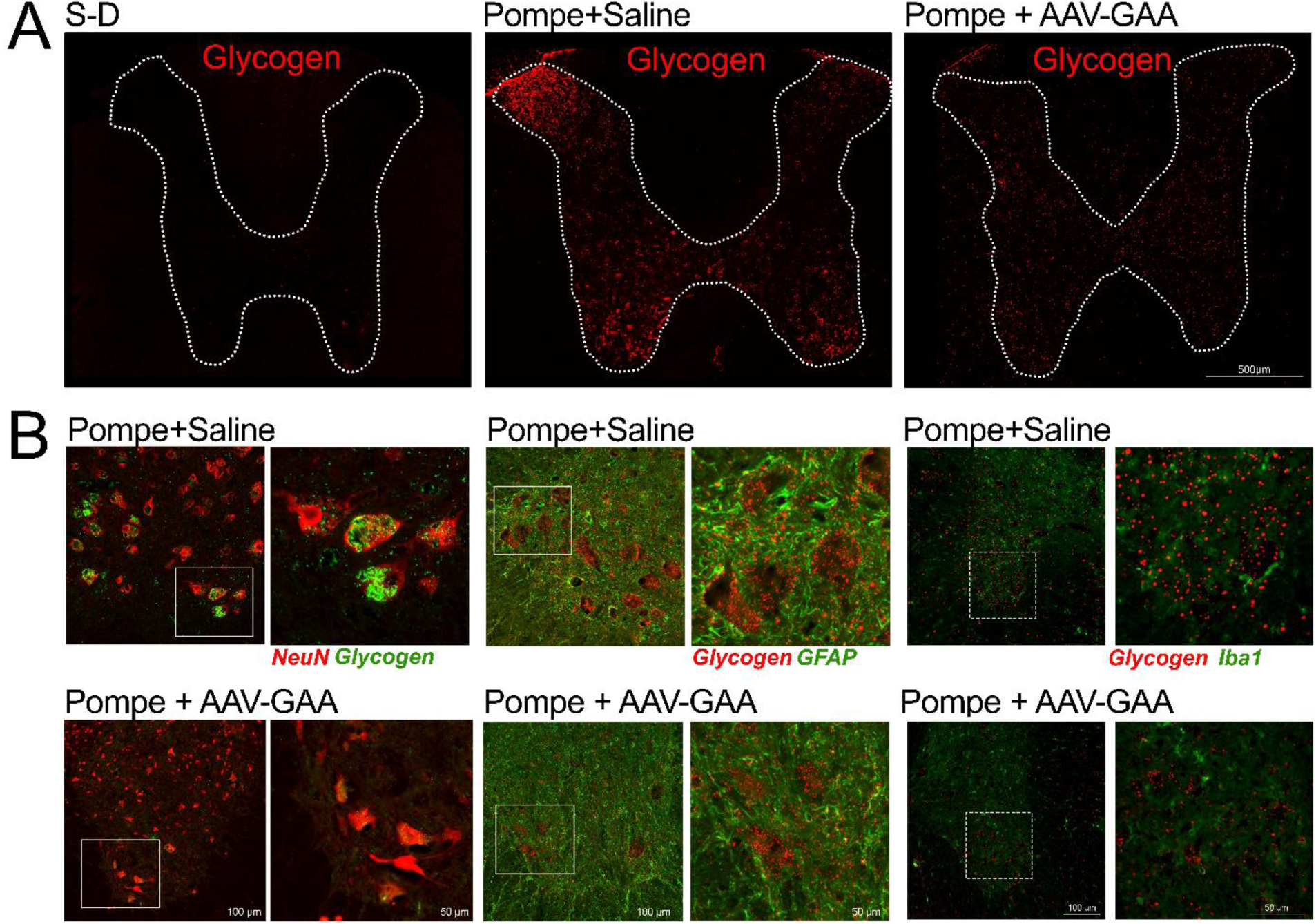
Representative photomicrographs of spinal cord tissue. Mid-cervical (C4-5) spinal cord sections were stained with NeuN (neurons) IV-58 (Glycogen), GFAP, and Iba1, and evaluated using fluorescence microscopy. The images demonstrate the expected marked increase in neuronal glycogen in Pompe+Saline rats, and a reduction in glycogen after AAV-GAA treatment. **A:** Low power images showing glycogen staining in spinal grey matter. **B-D:** Higher power images showing staining for neurons (NeuN) and glycogen (B), GFAP and glycogen (C), and Iba1 and glycogen (D). S-D: Sprague-Dawley

Assays to evaluate GAA activity and overall glycogen content in the heart and diaphragm were completed in a cohort of rats at age 6 months. As shown in **Figure 6A**, AAV-GAA treatment produced a substantial increase in GAA activity, with a corresponding reduction of glycogen in both tissues. The GAA activity assay was repeated at age 12 months, with demonstration of sustained increases in GAA activity in the heart, diaphragm, quadriceps, and spinal cord (**Figure 6B**). Biodistribution studies confirmed AAV vector genome detection in the central nervous system and diaphragm, but with highest levels expression in liver and heart (**Figure S2**).

**Figure 6.**
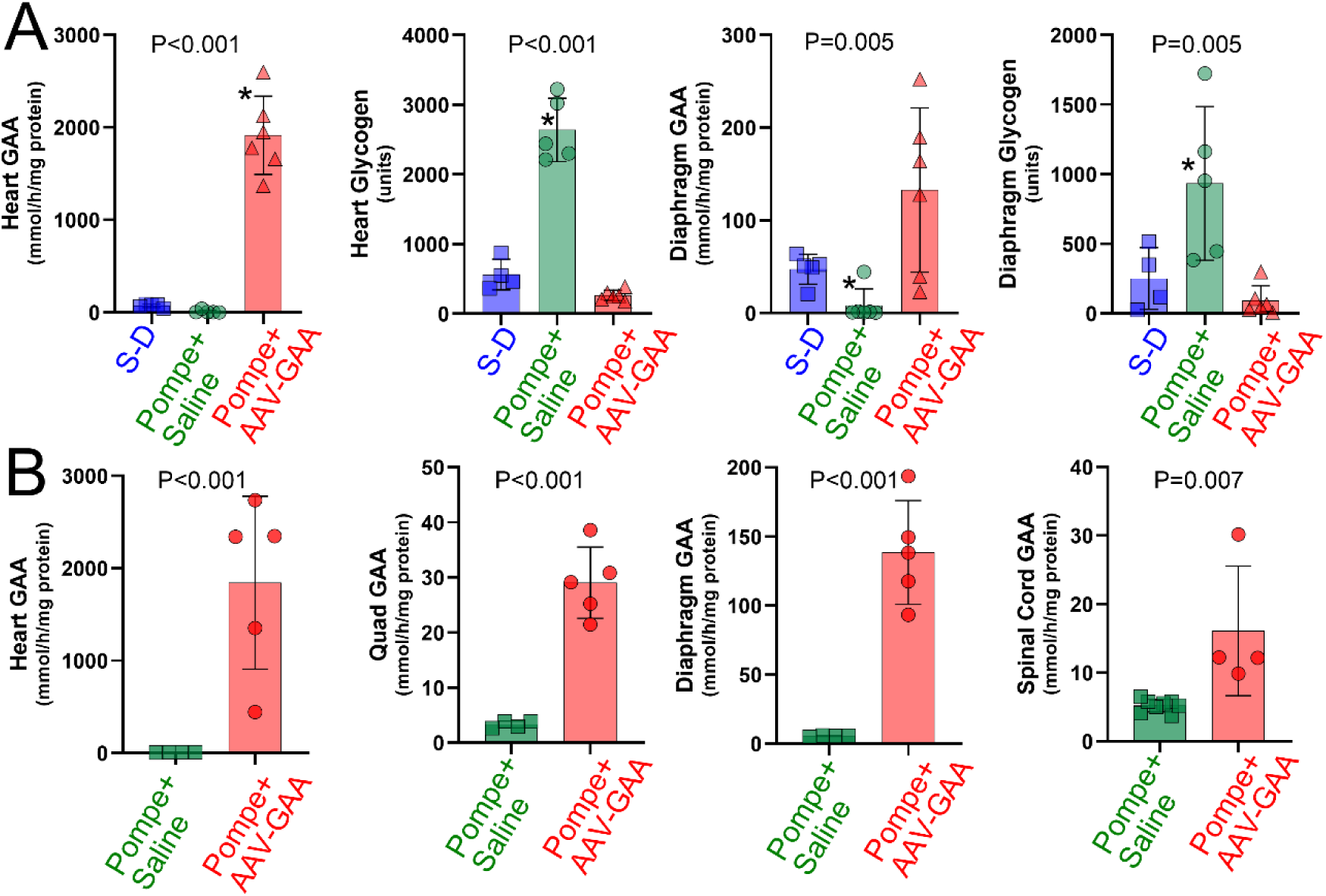
GAA activity and glycogen content. A: Assays done at age 6 months, neonatal AAV-GAA treatment increased GAA activity and reduced glycogen in heart and diaphragm. **B:** Assays done at age 12 month; neonatal AAV-GAA treatment increased GAA activity in heart, diaphragm, quadriceps and spinal cord. Statistical tests: A: 1-way ANOVA. *, different than Pompe+AAV-GAA. B: t-test. S-D: Sprague- Dawley

A comprehensive cardiac evaluation using MRI and ECG was performed in a cohort of rats at age 6 mo. Representative short-axis cardiac MRI images are shown in **Figure 7A**. *Ex vivo* assessment of the heart weight to body weight ratio (HW:BW) is shown in **Figure 7B**. *Gaa^-/-^* rats exhibited cardiomegaly with a 52% increase in HW:BW vs. Sprague-Dawley rats. However, the HW:BW ratio was normalized in *Gaa^-/-^* rats treated with AAV-GAA (**Figure 7B**). The MRI data were used to calculate cardiac volumes, and cardiac output was increased after AAV-GAA treatment (**Figure 7C**). Stroke volume (**Figure 7D**) and ejection fraction (**Figure 7E**) were variable in untreated Pompe rats but became much more consistent after AAV-GAA treatment. End-systolic and end-diastolic volumes (**Figure 7F**) were normalized in *Gaa^-/-^* rats following the AAV-GAA treatment, as was volume index (**Figure 7G**).

**Figure 7.**
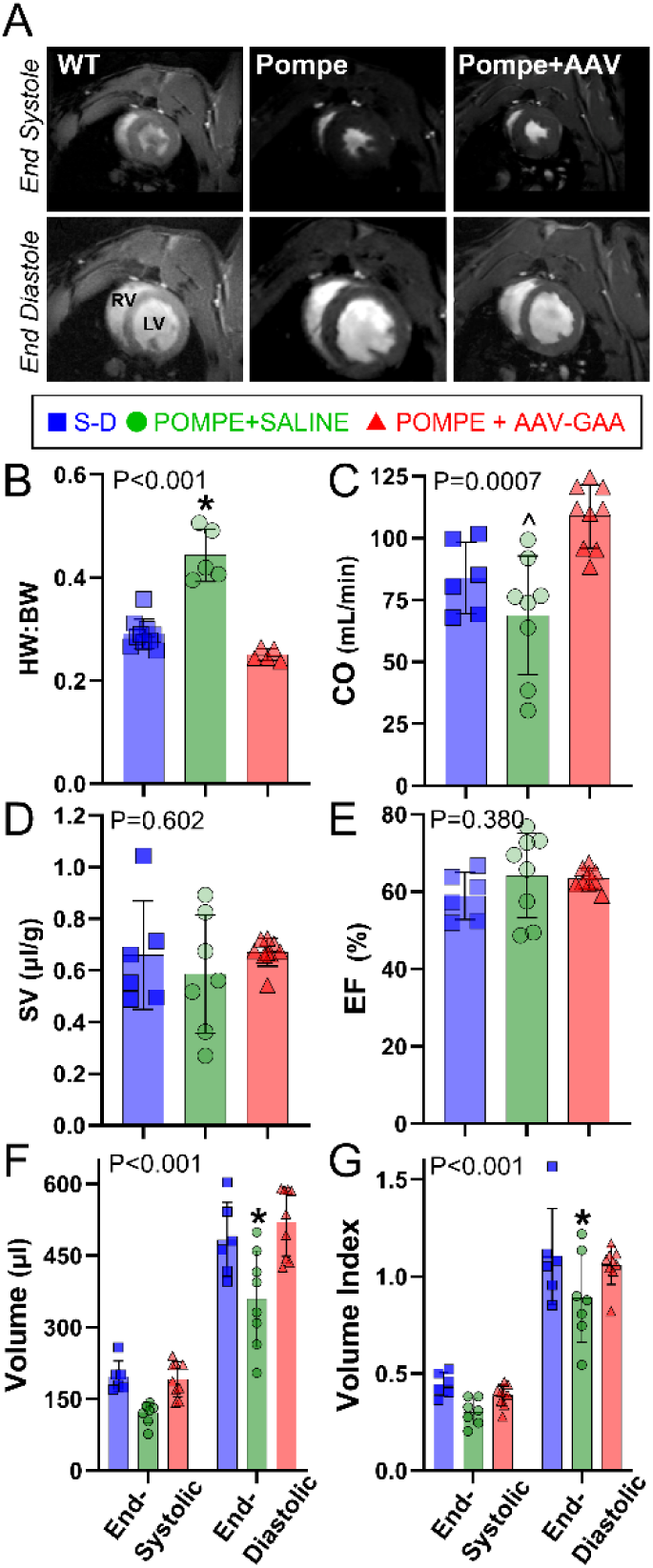
Impact of AAV-GAA on the heart. Representative MRI images are shown in panel **A**. **B:** *Ex vivo* assessment of the heart weight to body weight ratio (HW:BW) shows a reduction in size after AAV- GAA. **C:** Cardiac output (CO) was increased after AAV-GAA treatment. **D-E:** Stroke volume (SV) and ejection fraction (EF) are variable in saline treated Pompe rats but are more consistent after AAV-GAA treatment. **F:** End-systolic and end-diastolic volumes are normalized following the AAV-GAA treatment. **G:** Volume index is increased after AAV-GAA. *, p<0.05 vs. other two groups; ^, p<0.05 vs. Pompe+AAV- GAA. S-D: Sprague-Dawley

ECG was recorded using a 5-lead telemetry system; a summary of these measurements is shown in **Figure 8**. The R-R interval and R-wave amplitude were normalized in *Gaa^-/-^* rats that received the AAV-GAA therapy. Glycogen levels in the cardiac conduction system is one of the most sensitive measures of glycogenosis in IOPD.

**Figure 8.**
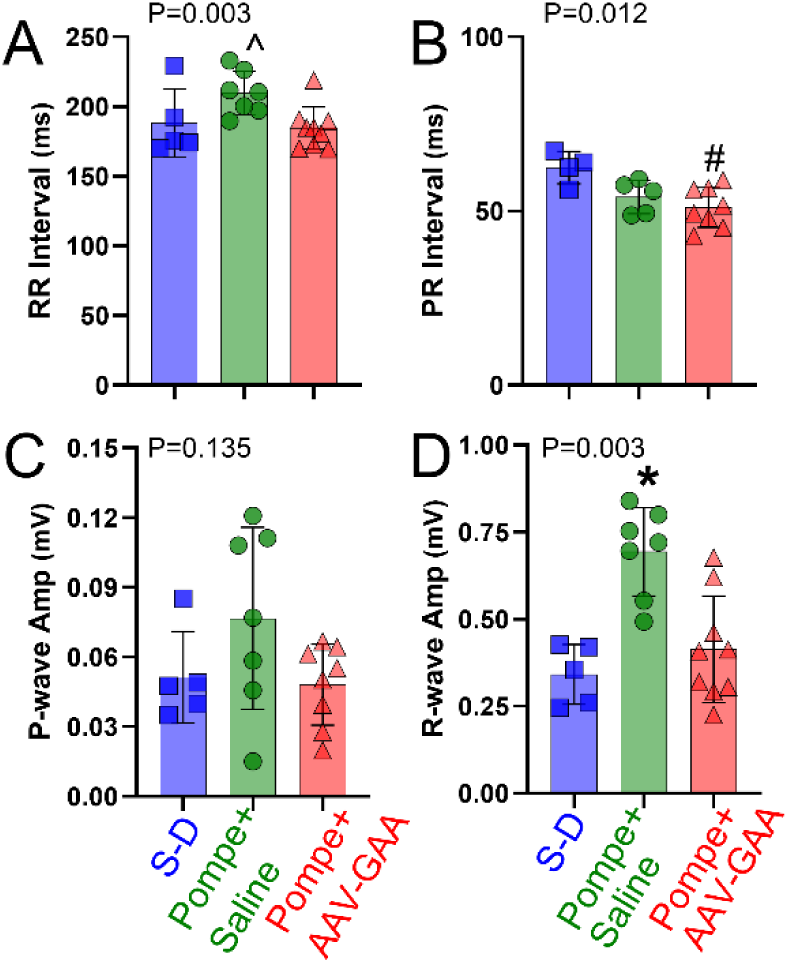
Impact of AAV-GAA on ECG. A: R-R interval, **B:** PR interval, **C:** P-wave amplitude, **D:** R- wave amplitude. The R-R interval and R-wave amplitude were normalized after AAV-GAA therapy. Statistical test: 1-way ANOVA. The treatment effect P-value is reported on each plot. *, p<0.05 vs. other two groups; ^, p<0.05 vs. Pompe+AAV-GAA; #, p<0.05 vs. Sprague-Dawley (SD)

Additional experiments provided an unbiased genome-wide screening of mRNA expression in the diaphragm and heart in *Gaa^-/-^* rats at 6 months age. This work was done as part of the validation of the rat model and did not include an AAV treated cohort. The results highlight the wide-ranging impact of GAA deletion and included to provide insight regarding the pathways that are impacted in Pompe disease. The GO ontology enrichment analysis is summarized in **Tables S1-S4**. Down-regulated genes in the *Gaa^-/-^*diaphragm indicate a marked decrease in the expression of genes involved in mitochondrial function, including those associated with the tricarboxylic acid (TCA) cycle, fatty acid metabolism. The GO analysis for up-regulated genes in the *Gaa^-/-^* diaphragm indicated activation of endoplasmic reticulum (ER)-associated pathways, including protein processing and the response to ER stress. Genes related to the immune response were also strongly upregulated. GO analysis for down-regulated genes in the *Gaa^-/-^* heart showed significant enrichment in ion transport pathways, specifically potassium ion transport and transmembrane channel activities. Pathways associated with fatty acid metabolism and lipid modifications were also down-regulated, consistent with alterations in lipid utilization and energy production. Analyses of up-regulated genes in the *Gaa^-/-^* heart revealed enrichment in immune-related processes, particularly innate immune response pathways, as well as extracellular matrix organization pathways. Additionally, the activation of metabolic processes, including those related to organic compound response and insulin-like growth factor signaling, points to metabolic stress and adaptations in the *Gaa^-/-^* heart.

## DISCUSSION

Our findings demonstrate that neonatal systemic treatment with AAV9-GAA leads to sustained correction of cardiorespiratory function in the Pompe rat model. Notably, the diaphragm muscle showed near-complete correction, as determined by spatial metabolomics and histological analysis. Whole-body plethysmography and phrenic nerve recordings further indicated improvements in respiratory function. Additionally, cardiac function was dramatically improved following gene therapy as shown by MRI and electrophysiological assessments. Collectively, these results contribute to the growing body of evidence supporting the efficacy of gene therapy in treating the most severe form of early onset Pompe disease. ^19,27–29^

### Cardiorespiratory failure in Pompe disease and the Gaa^-/-^ rat model

Early-onset Pompe disease occurs when functional GAA protein is absent or reduced, and this results in progressive hypertrophic cardiomyopathy^30^. Respiratory insufficiency is a hallmark of early-onset Pompe disease^4,31^ and can often be one of the first symptoms observed^30^. Both cardiac and respiratory impairments contribute to cardiorespiratory failure, which is the leading cause of mortality in the early-onset Pompe patients^30^.

The *Gaa*-null rat model described here closely mirrors many of the cardiorespiratory dysfunctions seen in IOPD disease. Cardiac MRI and post-mortem assessments confirmed the presence of cardiomegaly, with reduced cardiac volumes and ECG abnormalities compared to the unaffected control Sprague-Dawley rat. Whole-body plethysmography recordings in awake *Gaa^-/-^* rats revealed altered breathing patterns including reduced inspiratory tidal volume. There was also an indication of hypoventilation, which is consistent with the progressive respiratory failure seen in Pompe patients. ^32^ Additionally, the *Gaa^-/-^* rat displayed the prototypical histopathological changes in spinal motoneurons^5,8^ and the diaphragm^33^, as commonly observed in Pompe disease.

The majority of preclinical gene therapy and/or histopathology studies of Pompe disease have utilized the *Gaa*-null mouse model created by Raben.^34^ This model has been foundational for understanding disease mechanisms^6–8^ and advancing gene therapy treatments^16–18,35^. There is one prior report of a *Gaa*-null Pompe rat model, which similarly displayed glycogen accumulation, cardiomegaly, and reduced body weight.^36^ In that study, the Pompe rats showed early mortality, with death occurring by age 8 months of age, which is earlier than in Pompe mouse models^8^. In the current report, Pompe rats also showed early mortality, with 50% of rats surviving to 10 months of age (Figure S3). Overall, the data reported here are consistent with previous findings^36^, further establishing the *Gaa^-/-^* rat as a valuable model for studying the pathophysiology and treatment of Pompe disease. Further, mRNA gene array data (e.g., Tables S1-4) provide a hypothesis-generating resource for exploring the molecular mechanisms underlying neuromuscular decline in the absence of GAA activity. In the diaphragm, up-regulated genes were enriched in ER stress and immune response pathways, while down-regulated genes indicate compromised mitochondrial function and disrupted energy metabolism. In the heart, up-regulated genes suggest immune activation and extracellular matrix remodeling, while down-regulated genes highlight dysregulation in ion transport and lipid metabolism. These results collectively indicate distinct pathophysiological changes in muscle and cardiac tissues in response to Pompe disease, potentially driving dysfunction in a tissue-specific manner.

### Early-life AAV therapy in Pompe disease

An accumulation of data from animal models and initial clinical trials supports^14,37^ the use of gene therapy approaches in Pompe disease. While further optimization of viral vectors and immune management^19,28,29^ is required, momentum is growing for clinical gene therapy treatments in Pompe patients. With the expansion of newborn screening for Pompe disease in developed countries ^38–40^, the possibility of initiating gene therapy treatments at very early stages of disease is becoming increasingly feasible, as demonstrated in the current study.

Here, we utilized early-life systemic AAV9-GAA treatment to achieve widespread GAA activity and correct cardiorespiratory function. Prior studies have shown that intramuscular AAV9-GAA administration can effectively correct pathology in targeted skeletal muscles.^17,41^ Further, the retrograde movement of the viral vector following an intramuscular delivery can drive robust GAA expression in motor neurons^17^. Thus, intramuscular AAV delivery offers a powerful strategy for targeting gene expression across the entire motor unit (*i.e*., both myofibers and motor neurons). This method is highly effective at targeting lingual motor units in Pompe models.^17,41^ However, to fully prevent cardiorespiratory decline, systemic gene therapy treatments capable of targeting the heart, skeletal muscles, and CNS will be required.^19^

The recent study by Munoz et al. treated 4-week-old Pompe rats with an AAV-GAA vector incorporating an optimized muscle-specific promoter and a transcriptional cis-regulatory element.^36^ Following systemic delivery via tail vein, muscle glycogen levels, mass, and strength were normalized when evaluated at 20 weeks of age (i.e., 4-mo post-treatment). Thus, this approach was highly effective in correcting both skeletal and cardiac pathology. A clinical concern, however, is that correcting muscle function without addressing CNS pathology is likely to lead to the emergence of a neural phenotype.^42^ Additionally, data from Pompe patients ^10,11,13^ and animal models^8,43^ indicate that the neural control of breathing progressively worsens with aging, likely due to prominent pathology in respiratory motoneurons ^5,7^. To address this issue, Keeler et al. aimed to target both muscle and the CNS by treating Pompe mice with an AAV vector (AAVB1) that has high tropism for skeletal muscle and the CNS ^18^ Three-month-old Pompe (*Gaa^-/-^)* mice were systemically treated with AAVB1 encoding GAA, which led to GAA expression in the heart and diaphragm, resulting in sustained improvements in respiratory function.

Building on prior work, the current experiments make a few key advances. First, we demonstrate sustained correction of cardiac size and ECG activity in Gaa-null rats. This was accompanied by correction of cardiac function following AAV-GAA treatment, as shown by a comprehensive evaluation using MRI. Second, we used spatial metabolomic methods ^44^ to demonstrate the remarkable impact of AAV-GAA on the diaphragm muscle. Specifically, diaphragm glycans were normalized post-gene therapy, which is fundamentally important in a glycogen metabolic disorder such as Pompe. Finally, gene therapy-treated rats showed a normalized breathing pattern across development (i.e. 6-12 mo.). Both inspiratory tidal volume and minute ventilation were considerably increased in AAV-GAA-treated rats compared to saline treated Pompe rats. This improvement was further supported by neurophysiological recordings of the phrenic nerve and histological evaluation of neuron morphology, suggesting a positive impact of GAA replacement on respiratory neural control circuits.

## Conclusion

We conclude that early-life treatment with an AAV9 vector driving GAA expression can lead to sustained cardiorespiratory correction in a Pompe rat model. This adds to the body of work supporting the potential of gene therapy for IOPD disease ^19,28,29^. With the expansion of newborn screening for Pompe disease, early detection and intervention become feasible, making early-life therapy a possibility^1^. However, challenges remain, particularly in managing immune responses ^19,28^. In the current studies, AAV- GAA was delivered before rat immune system had fully matured, eliminating the need for immunosuppression. Immune responses to the transgene product (i.e., GAA) will be of particular concern for the cross-reactive immunologic material (CRIM)-negative early-onset Pompe patient. Nevertheless, continued development of gene therapy strategies offers high potential for more stable and widespread expression of GAA to address both systemic and CNS manifestations of the disease compared to current approaches.

## METHODS

Procedures were approved by the Institutional Animal Care and Use Committee at the University of Florida. Rats were housed with littermates under temperature-controlled conditions with 12-hr light/dark cycles and food and water *ad libitum*.

### Gaa^-/-^ rat model

Creation of the rat model is summarized in **Figure S4**. Zinc finger nucleases (ZFN) are DNA binding proteins that enable targeted genomic editing by producing double stranded DNA cuts. The design and validation of ZFN reagents (Millipore Sigma, CompoZr^®^ Custom ZFN) and procedures used to create a Pompe rat model by targeting the acid alpha glucosidase (*Gaa*) transgene were performed as previously described.^45–47^ The ZFN binding (shown in CAPS) and cutting (shown in lowercase) sequence design is CACTGCCCTCCCAGCacatcACAGGCCTGGGTGAG. The ZFNs were delivered to Sprague Dawley (SD) embryos which were then implanted in pseudo-pregnant SD females.^48^ Initial screening of ZFN-modified progenies identified a series of disruption to the *Gaa* coding sequence. Founder animals were selected based on Sanger sequencing of PCR amplicons (ICBR core, University of Florida) that identified a number of animals containing identical deleted sequences within the *Gaa* open reading frame (designated *Gaa^-/-^)*. Male *Gaa^-/-^* KO x Female SD rats were paired and subsequent offspring were bred to heterozygosity or homozygosity for establishment of the Pompe KO rat colony. *Gaa^-/-^* rats were sequenced to confirm *Gaa* deletion.

The present study did not directly test the impact of the AAV-GAA therapy on lifespan in the *Gaa^-/-^* rat model. The study design necessitated tissue collection or physiological measurements at set endpoints. In **Figure S3** we present the survival curve obtained from our *Gaa^-/-^*rat colony over the last several years. Compared to the well-established expected lifespan of the male Sprague-Dawley rat (*e.g*., approximately 2 years^49^) the *Gaa^-/-^*rat exhibits early mortality.

### Study design

Male rats were used because respiratory decline has been reported to be more severe in males with Pompe disease^50^. Further, our prior report in *Gaa^-/-^* mice^8^ and initial data in rats (**Figure S5**) indicated that the respiratory phenotype in the *Gaa^-/-^* rat model is more severe in males, which is consistent with clinical reports that respiratory decline progresses faster in males^50^. Pompe rats were treated with AAV- GAA (see next section) or sham (saline of equal volume) on post-natal day 1. Separate cohorts were evaluated for cardiac function using *in vivo* MRI (age 5-6 mo.) and respiratory function using whole body plethysmography in awake rats (age 6-12 mo.) and neurophysiological recordings of the phrenic nerve under anesthesia (age 5-6 mo.). Sample sizes are reported with the description of the methods for each outcome measure. Tissues were harvested at 6-12 months for histological, molecular and MALDI assessment.

### AAV

Single stranded AAV9 vectors encoding the human GAA (hGAA) protein, driven by the desmin promoter (AAV9-Des-hGAA) were used. The vector was packaged, purified, and titered at the University of Florida Powell Gene Therapy Center Vector Core Laboratory. Vectors were purified by iodixanol gradient centrifugation and anion-exchange chromatography as previously described ^51^. A single intravenous injection of 5e13vg/kg AAV9-Desmin-hGAA was delivered into the temporal facial vein at postnatal day 0 (P0). Briefly, rat pups were cryo-anesthetized for ∼1 minute. The vector was injected using a 30G tuberculin syringe in a maximal volume of 20ul. Animals were immediately returned to their home cage and monitored for recovery following the procedure.

### Magnetic resonance imaging (MRI) and electrocardiography

For cardiac MRI, we studied n=6 SD, n=8 Pompe + saline, and n=9 Pompe + AAV-GAA. A 4.79T Bruker Avance spectrometer (Bruker BioSpin Corporation, Billerica, MA) at the University of Florida AMRIS facility was used as previously described^20^. Rats were anesthetized (1.5% isoflurane and 1L/min. oxygen) and positioned on a quadrature transmit- and-receive surface coil. Single short-axis slices were visualized along the left ventricle. Images were acquired using IntraGate and processed using CAAS MRV 4.3 (Pie Medical Imaging, Maastricht, The Netherlands) throughout the complete heartbeat cycle. Images at end systole and end diastole were analyzed to obtain systolic volume (SV), cardiac output (CO), ejection fraction (EF), end systolic volume (ESV), end diastolic volume (ESV), end systole (ES) and end diastole (ED) mass.

Electrocardiography (ECG) recordings were performed as previously described^20^ at 3 and 5.5-6 months of age in SD (n=5), Pompe+saline (n=7) and Pompe+AAV-GAA (n=9). Briefly, rats were anesthetized using 1.5% isoflurane and 1L/minute oxygen. Five electrode leads were placed in the tail, left lower leg, right scapular region, right forelimb, and left forelimb to acquire steady ECG tracings using AD Instrument Chart software. The Q amplitude, R amplitude, RR interval, and PR interval were averaged over a period of 3min. and reported as a single value for each rat.

### Plethysmography

A flow-through whole body plethysmograph was used to measure overall ventilation in awake, freely moving animals as described previously^52^. A CO_2_ sensor was at the outflow point enabled measurement of metabolic CO_2_ production (V̇CO_2_) using Fick’s principle ^53^. The experimental protocol consisted of a 45-minute baseline under normoxic air (21% O_2_, 79% N_2_), followed by 7-min hypoxic (10.5% O_2_ balance N_2_) period, followed by a 20-min post-hypoxia recording period under normoxic air, finally ending with a 7-minute maximal chemoreceptor challenge (10.5% O_2_, 7% CO_2_ balance N_2_). Rectal temperature was assessed at the end of all recording sessions. Tidal volume was calculated using the equations developed by Drorbaugh and Fenn ^54^.

### mRNA gene array

An mRNA gene array evaluation was completed for the heart and diaphragm in group of age 5 mo. male *Gaa^-/-^* (n=3) and Sprague-Dawley rats (n=3) with no prior treatment. This analysis was conducted to describe the new *Gaa^-/-^* rat model and to provide a hypothesis generating data set for future work. We did not have a specific *a priori* hypothesis regarding the transcriptome data, which are summarized in **Tables S1-4.** The complete RNAseq datasets are available at https://odc-sci.org/ under “David Fuller Laboratory”.

Rats were injected intraperitoneally with Beuthanasia^®^ (150 mg/kg) solution. Tissues were harvested, placed into RNA Later (Life Technologies, Carlsbad, CA, USA), and stored at -80°C. RNA extraction was performed using TRIzol and isolated total RNA was purified using a RNeasy Mini kit (Qiagen, Valencia, CA). The resulting quantity and purity of total RNA was tested through absorbance spectrophotometry at 230, 260 and 280 nm. RNA samples were sent to the Boston University Medical Center Microarray Core Facility for analysis using the Affymetrix Rat Gene Array 2.0ST. Gene expression profiles were analyzed using the Rat Transcriptome Assay 1.0 microarray platform. Twelve samples were included in the study, with the following groups: heart (n=3) and diaphragm tissue (n=3) from Sprague-Dawley rats, and heart (n=3) And diaphragm (n=3) from Pompe rats (n=3). Microarray data were processed using the Affymetrix Expression Console to obtain log2-transformed gene-level expression values, followed by normalization using the Robust Multiarray Average (RMA) method. Data quality was assessed using Relative Log Expression (RLE) and Area Under the Receiver Operating Characteristics Curve (AUC) values, which were all above 0.8, indicating high-quality data.

Differential expression analysis was conducted using a linear modeling approach. Two linear models were used to assess the effects of disease status (Pompe vs. Sprague-Dawley) and tissue type (diaphragm vs. heart) on gene expression: 1) main effects model: expression ∼ disease + tissue; 2) interaction model: expression ∼ disease + tissue + disease:tissue. Student’s two-sample t-tests were performed for each effect, and Benjamini-Hochberg False Discovery Rate (FDR) correction was applied to control for multiple hypothesis testing. Results were considered significant at an FDR-corrected p-value (q value) threshold of <0.05. Probesets with low overall expression were filtered out to reduce the likelihood of false positives.

Differentially expressed genes were subjected to Gene Ontology (GO) analysis using the DAVID Bioinformatics Resource. Functional enrichment was assessed to identify overrepresented biological processes, cellular components, and molecular functions associated with the altered gene expression patterns. Pathways were considered significantly enriched at a p-value threshold of <0.05 after multiple testing correction. All statistical analyses and visualizations were performed using the R environment for statistical computing (version 2.15.1), and differential expression analysis utilized the limma package (version 3.14.4). Gene ontology enrichment was conducted using DAVID (Sherman et al., 2022).

### Phrenic nerve recordings

Details of the surgical methods have been described ^55^. Anesthesia was initially induced with 3% isoflurane and then maintained with 3% isoflurane, 65% O_2_, 1% CO_2_ mixture delivered via a nose cone. After demonstration of loss of pedal withdrawal and corneal reflexes rats were tracheotomized, vagotomized and ventilated (VentElite, model 55-7040; Harvard Apparatus Inc.) at 65-75 breaths/min and tidal volume of 7 mL/kg. Urethane anesthesia was induced via femoral vein infusion (1.7 g/kg, 6 mL/h) followed by a continuous infusion of 8.4% sodium bicarbonate and lactated Ringer’s (2 mL/h). Pancuronium bromide was used to induce neuromuscular blockade (3 mg/kg, IV Sigma-Aldrich, St Louis). Arterial blood pressure and partial pressure of CO_2_ (PaCO_2_), O_2_ (PaO_2_), and pH (ABL 90 Flex, Radiometer, Copenhagen, Denmark) were measured via a femoral catheter. The right phrenic nerve was recorded using a glass suction electrode filled with saline. Signals were amplified (Model 1700, A-M systems, Everett, WA), band-pass filtered (100 Hz–3kHz), digitized (16 bit, 25,000 samples/channel, Power 1401, CED), and integrated (∫) with a time constant of 0.05 sec using Spike2 software (Cambridge Electronic Design, UK). The CO_2_ apneic threshold for phrenic bursting was determined, baseline recordings were made for 15 min, and rats were exposed to acute hypoxia (11.5% O_2_) as described ^55^. Spike2 software was used to record data (version 10.01, Cambridge Electronic Design). Data were analyzed using a custom MATLAB code (MathWorks, R2019a)^55^.

### Vector biodistribution

The biodistribution of the AAV9-Desmin-hGAA vector was analyzed using real-time PCR detection as previously described ^20,35^. Data are expressed as vector genome/diploid genome (VG/dp).

### Immunohistochemistry

Animals (WT rats (n=3), *Gaa^-/-^* rats (n=3), and *Gaa^-/-^*rats (n=3) injected with AAV9-Desmin-hGAA) were deeply anesthetized, euthanized by exsanguination, and transcardially perfused with chilled 4% paraformaldehyde in 0.1 M phosphate buffered saline (PBS, pH=7.4). The spinal cord was resected from brainstem to lower lumbar sections, postfixed in 4% paraformaldehyde for 24 h, and transferred to 30% sucrose in 0.1 M PBS (pH 7.4) for 3 days at 4°C. Brainstem and spinal cords were subsequently embedded in cryomolds (VWR, Radnor, PA), sectioned at 20 μm and thaw mounted directly onto slides.

Methods for visualizing GAA expression have been described ^17^.Tissues were incubated overnight in primary antibody (1:2000 rabbit polyclonal GAA antibody, Covance, Emeryville, CA), washed with PBS, incubated in a biotinylated anti-rabbit IgG secondary antibody (1:200 Vector Laboratories, Burlingame, CA), and treated with DAB for visualization with bright field microscopy.

Another set of slides were processed with primary antibodies against IBA1 (1:500; Wako # 019-19741), GFAP (1:500; Encor #MCA-1B7), or NeuN (1:500; Encor #MCA-1B7). For immunohistochemistry, sections were blocked (10% serum, 60 min), blocked in primary antibodies overnight at 4°. After primary incubation and three serial washes with 1 × PBS, secondary antibodies were incubated for two hours at room temperature. Secondary antibodies were washed off with 1 × PBS and coverslips were mounted with Vectashield hardset mountant (Vector Laboratories). Tissue sections were imaged and stitched using a 10x and 20x objective on a Keyence microscope (BZ-X700, Keyence Corporation of America, Itasca, IL).

### GAA and glycogen assays

The heart, diaphragm, quadriceps and spinal cord were analyzed for GAA activity as previously described ^41,56^. Tissues were harvested, flash frozen in liquid nitrogen, and maintained at -80°C until biochemical analyses were performed. Tissues were homogenized in water containing a protease inhibitor cocktail and exposed to three freeze-thaw cycles. Homogenates were subsequently centrifuged at 4°C, the supernatant was incubated for 1 hour at 37°C, and GAA activity as low as 0.05µmol/l/h*µg was assessed by measuring the cleavage of 4-methylumbelliferyl-α-D-glucopyranoside. Measurement of glycogen content in cardiac and diaphragm was done using the Glycogen Assay Kit (ab65620; Abcam, Cambridge, MA), following the manufacturer’s instructions as in our prior publication^56^.

### MALDI-MSI

To map metabolic alterations in the diaphragm, we performed 2D MALDI imaging^44,57^ of the metabolome, lipidome, and glycogen on serial diaphragm sections from WT, GAA, and GAA-AAV rats. Diaphragm tissues were frozen immediately post-dissection and sectioned at 10 µm thickness. Each section underwent MALDI mass spectrometry imaging (MALDI-MSI) with specific matrices tailored for metabolite and lipid mapping. We applied the N-(1-Naphthyl) ethylenediamine dihydrochloride (NEDC) matrix for initial metabolome and lipidome scans with a spatial resolution of 10 µm, utilizing a Bruker timsTOF flex MALDI-TOF instrument equipped with a Smartbeam2 laser for high precision. For glycogen and glycan imaging, tissue sections were treated with an enzymatic solution containing isoamylase and PNGase F to release these biomolecules. An HTX-M5 sprayer station (HTX Technologies) was used to coat each slide with 0.2 mL of this enzyme solution, containing 3 units of isoamylase and 20 mg of PNGase F per slide. The sprayer nozzle temperature was set to 45°C, with a spray velocity of 900 mm/min, ensuring even application. Slides were then incubated at 37°C in a humidified chamber for 2 hours, followed by desiccation to remove moisture before matrix application. For the matrix, we dissolved 0.04 g of α-cyano- 4-hydroxycinnamic acid (CHCA) in 5.7 mL of a 50% acetonitrile and 50% water solution, with an additional 5.7 μL of trifluoroacetic acid to enhance ionization. This CHCA solution was applied using the HTX-M5 sprayer, optimized for consistent coverage, thereby enabling high-resolution imaging of glycans and glycogen. MALDI images were acquired using the Bruker timsTOF flex MALDI-TOF instrument. The instrument was operated in reflectron mode for optimal resolution and high mass accuracy. Imaging data were collected and processed using Bruker SCiLS software, which facilitated control over imaging parameters and acquisition settings. The resulting images were exported for further analysis. The acquired MALDI-MSI data were then processed through the Sami pipeline for accurate slice alignment, providing a high-resolution view of metabolic distributions across the diaphragm. Metabolomics, lipidomics, and glycomic data were curated in MetaboAnalyst for hierarchical clustering and heatmap analysis, allowing visualization of metabolic patterns and identification of significant biochemical alterations among WT, GAA, and GAA-AAV groups.

### Muscle Fiber Typing

Immunohistochemical and imaging analysis was performed on diaphragm muscle cross sections. After animals were euthanized, the diaphragm was immediately harvested and flash frozen in liquid nitrogen. Sections were cut at 10um on a cryostat, mounted on slides, and air dried overnight at room temperature. Sections were fixed in ice-cold acetone for 10 minutes and subsequently air dried. Tissue was rehydrated for 5min in 1X PBS and incubated in Super Blocker (Pierce) for 40mins at room temperature. Slides were incubated O/N at 4°C using the following primary antibodies: Laminin (rabbit at 1:300, Sigma #L9393), MHC Type I A.4.840 (mouse at 1:30, Developmental Studies, IgM), and MHC Type IIa SC-71 (mouse at 1:25, Molecular Probes #A21121). The next day, slides were washed 2 x 5mins in 1X PBS and incubated at room temperature and in the dark for 1h with the following secondary antibodies: Alexa 405 anti-rabbit (1:250), Alexa 495 anti-mouse IgM (1:500), and Alexa 488 anti-mouse IgG (1:500). Slides were washed 2 x 5min in 1X PBS, placed in 4% paraformaldehyde for 3mins, washed 2x5min in 1X PBS, and cover slipped with Dako fluorescence mounting medium without DAPI. The slides were observed under a fluorescence microscope using the following filter settings: DAPI filter (blue) for Laminin, Texas Red filter (red) for Type I, and GFP filter (green) for Type IIa. Both diaphragm muscle cross sectional area and fiber type frequency were analyzed using Image J.

### Statistics

Statistical tests were conducted using GraphPad Prism software. Statistical significance was set at alpha level *p<0.05, and values in figures are reported as mean ± 1 standard deviation.

## Acknowledgments

AAV production by the University of Florida Powell Gene Therapy Center Vector Core Laboratory. Vector genome analysis by the University of Florida Toxicology Core. Cardiac imaging performed at the University of Florida Advanced Magnetic Resonance Imaging and Spectroscopy facility. Support: NIAMS K01AR066077 to DJF; R01HD052682 (DDF and BJB), R01AG066653 (RCS), R01CA266004 (RCS), R01AG078702 (RCS), RM1NS133593 (RCS).

## Author contributions

Conceptualization: DDF, RJF, BJB; Formal analysis: DDF, SR, PT, EB, AGT, DC, RAR, RCL, VC; Funding acquisition: DDF, DJF, BJB, RCS; Investigation: SR, PT, MP, AGT, LV, DC, RAR, RCL; VC; Methodology: DJF, AGT, LV, BJB; Project administration: SR, MP, LV; Resources: MSG; Supervision: DDF, DJF, BJB; Visualization: DDF, ESB, VNJ, RAR; Writing – original draft: DDF, VNJ; Writing – review & editing: DDF, RCS, DJF, MC, BJB

