## Supplemental File for "Neonatal systemic gene therapy restores cardiorespiratory function in a rat model of Pompe disease"

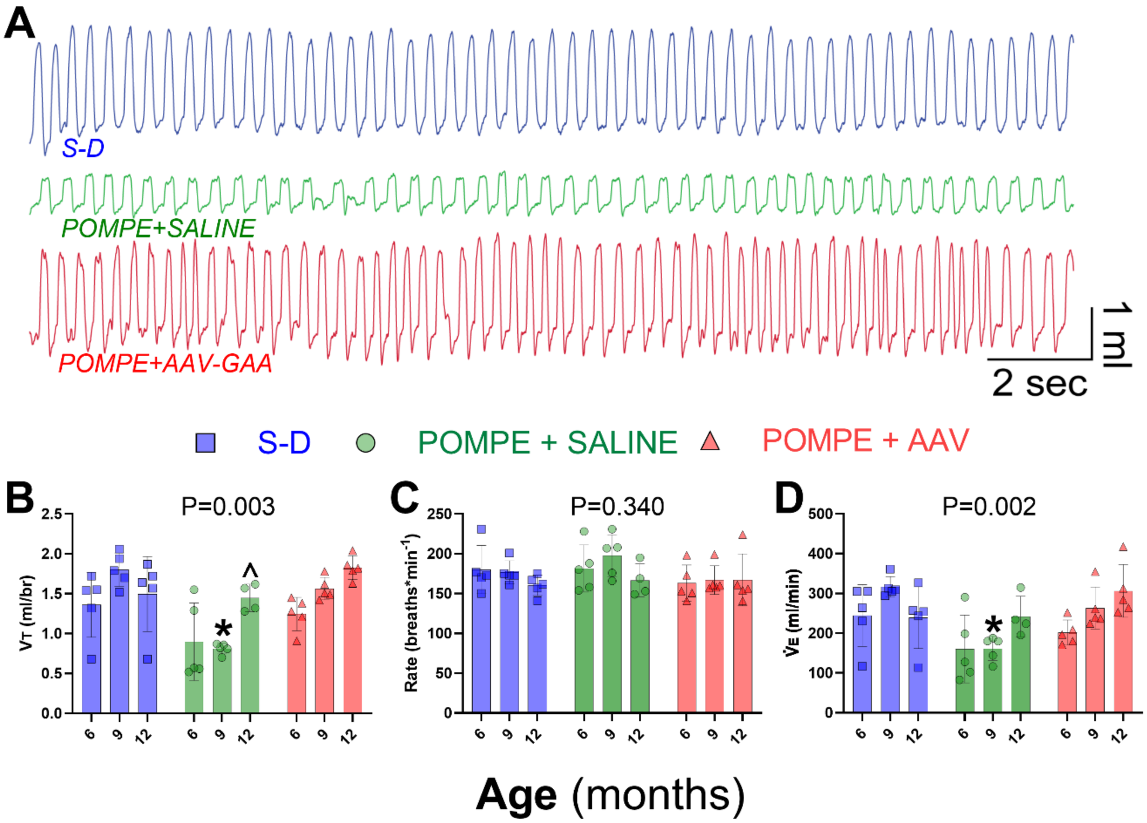

**Figure S1. Ventilation during an acute respiratory challenge.** Data obtained during a brief exposure to a 10% $O_2$, 7% $CO_2$ gas mixture. **A:** Example of breathing patterns measured using whole body plethysmography. **B:** Tidal volume (VT, ml/breath) is restored after AAV treatment. **C:** Respiratory rate is similar across the three groups. **D:** Minute ventilation ($\dot{V}$E, ml/min) is restored after AAV treatment. Statistical test: 2-way RM ANOVA. The treatment effect P-value is reported on each plot. *, $p<0.05$ *vs.* other two groups; ^, $p<0.05$ *vs*. Pompe+AAV-GAA. S-D: Sprague-Dawley

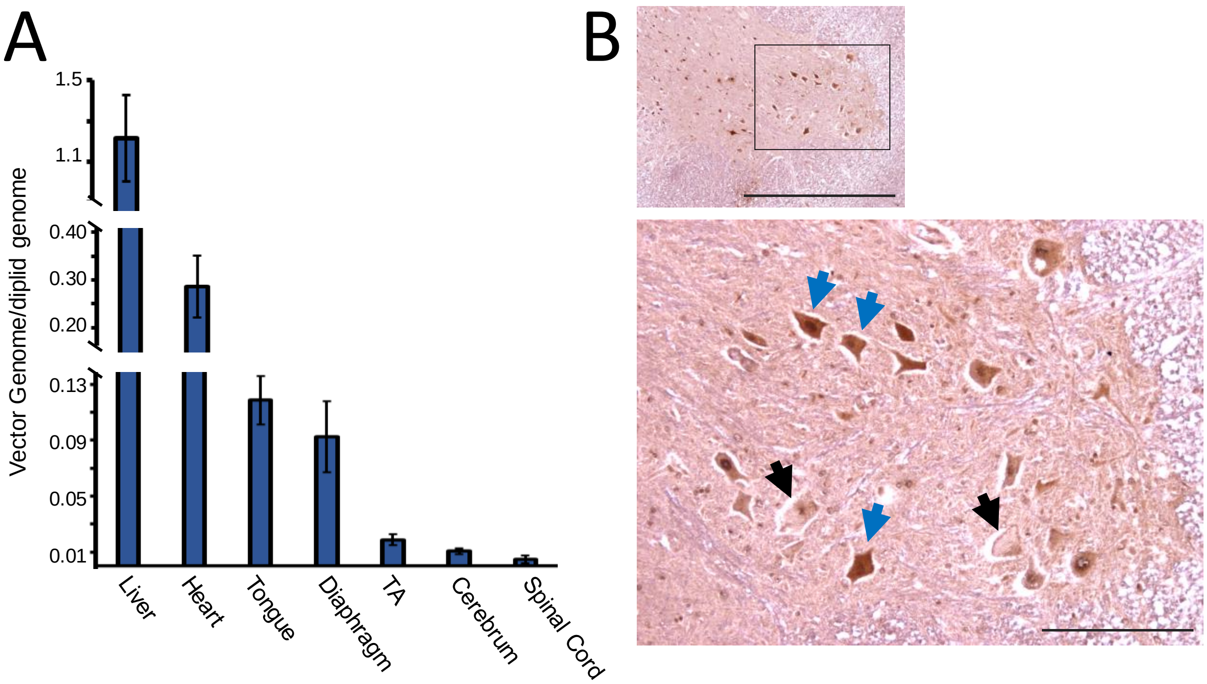

**Figure S2. Biodistribution of AAV9-Des-hGAA vector in AAV treated *Gaa*$^{-/-}$ rats and histological evidence of GAA expression. A.** The biodistribution of the AAV9-Desmin-hGAA vector was analyzed in 6mo old *Gaa*$^{-/-}$ rats that were treated with gene therapy at P0. The vector genome/diploid genome (VG/dp) is graphed for (left to right): the liver, heart, tongue, diaphragm, tibialis anterior (TA), cerebrum, and cervical spinal cord. **B.** Immunohistochemical example of GAA immunostaining in the mid-cervical (C4) spinal cord of an age 6-mo *Gaa*$^{-/-}$ rat that was treated with AAV9-Des-hGAA at P0. The example is included to illustrate the difference in histological appearance between GAA-positive motor neurons (dark brown stain; examples indicated by blue arrows) and GAA-negative motor neurons (examples indicated by black arrows). Note that GAA-positive cells are smaller and do not have the vacuolar appearance typical of GAA-negative cells. The bottom panel is a high magnification view of the area highlighted by the box in the top pane. Scale bars: 500 µm (top panel), 100 µm (bottom panel).

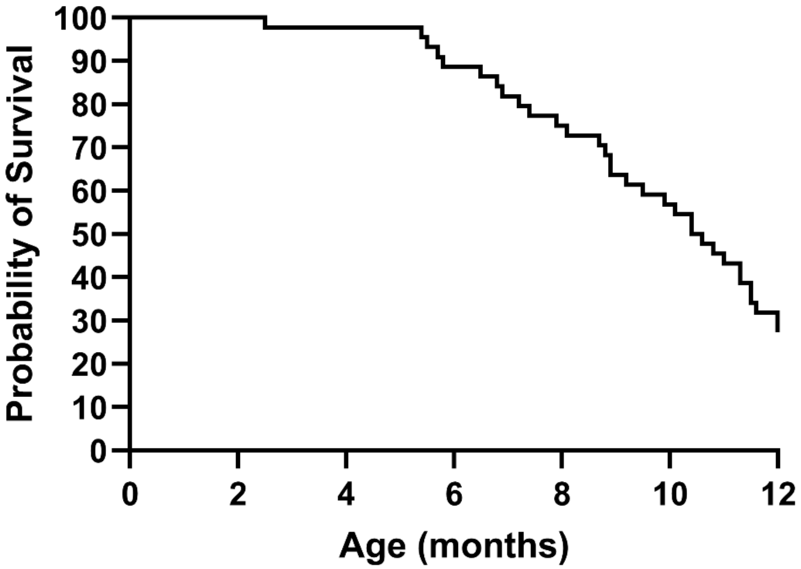

**Figure S3. Survival curve for male Pompe (*Gaa*$^{-/-}$) rats.** The survival data are for untreated male rats (n=44) in our colony over the last several years.

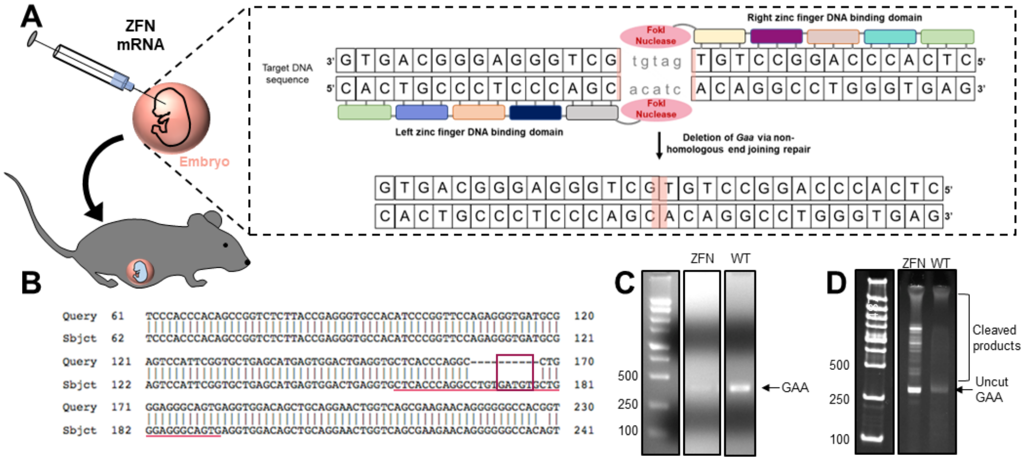

**Figure S4. Development and validation of the novel Zinc Finger Nuclease Pompe disease rat model.** **A:** Schematic of the development of the novel Pompe disease rat using Zinc Finger Nuclease (ZFN) gene editing technology. Boxed inset: The left and right zinc finger DNA binding domains bind to the target *Gaa* gene DNA sequence so the FokI nucleases can cleave $\geq$5 base pairs to cause mismatched base pairs via non-homologous end joining repair and knockout the *Gaa* gene. **B:** Example DNA sequencing readout of a successfully deleted *Gaa* gene in a founder ZFN rat. The spacer region that the FokI nucleases bind to (maroon box) and the target DNA sequence that are bound by the zinc finger nuclease binding domains (pink underline) are indicated.

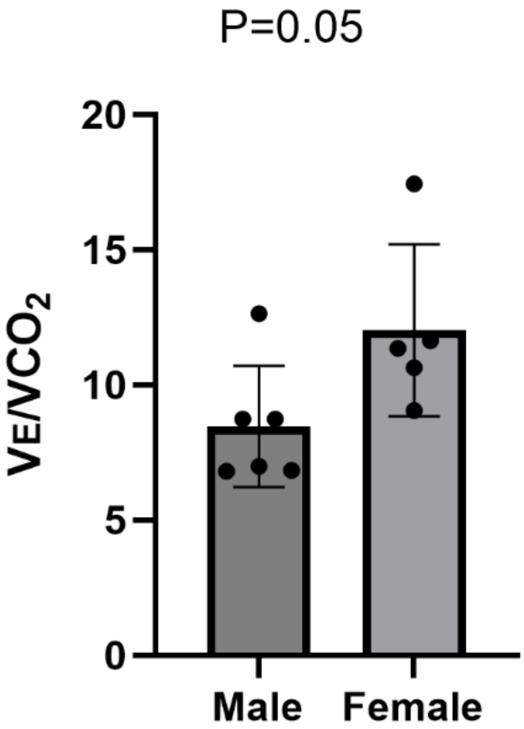

**Figure S5. The ratio of minute ventilation ($\dot{V}E$) to metabolic $CO_2$ production ($\dot{V}CO_2$) in male and female Pompe (*Gaa*$^{-/-}$) rats.** The reduction in the $\dot{V}E/\dot{V}CO_2$ ratio in males is consistent with a relative hypoventilation. Unpaired t-test.

**Table S1. GO ontology enrichment analysis for downregulated genes in the diaphragm of *Gaa*$^{-/-}$ rats.** The top three clusters are presented. Each row represents an enriched GO term or pathway, including details on the number and percentage of genes from the list associated with each term, enrichment significance (p-value), fold enrichment, and the statistical corrections applied (Bonferroni, Benjamini, and false discovery rate or FDR). % is the percentage of genes that are annotated to a specific GO term; "List.Total" stands for the total number of DE genes; "Pop.Hits" stands for the number of background population genes associated with the listed ID of gene ontology term; "Pop.Total" stands for the total number of background population genes. “Fold Enrichment” represents the ratio of genes from that belong to the specific GO term, compared to the expected proportion of genes annotated to that term in the whole genome.

| Category | Term | Count | % | P value | List Total | Pop Hits | Pop Total | Fold Enrichment | Bonferroni | Benjamini | FDR |
|---|---|---|---|---|---|---|---|---|---|---|---|
| ***Annotation Cluster 1*** | ***Enrichment Score: 31.35*** | | | | | | | | | | |
| GOTERM_CC_DIRECT | GO:0005739~mitochondrion | 141 | 23.4 | 4.99E-43 | 572 | 1460 | 21777 | 3.7 | 2.31E-40 | 2.31E-40 | 2.23E-40 |
| UP_KW_DOMAIN | KW-0809~Transit peptide | 76 | 12.6 | 2.81E-42 | 363 | 401 | 13371 | 7.0 | 7.30E-41 | 7.30E-41 | 7.30E-41 |
| UP_SEQ_FEATURE | TRANSIT:Mitochondrion | 65 | 10.8 | 1.49E-38 | 565 | 322 | 22265 | 8.0 | 1.72E-35 | 1.72E-35 | 1.70E-35 |
| UP_KW_CELLULAR_COMPONENT | KW-0496~Mitochondrion | 105 | 17.4 | 2.13E-37 | 402 | 962 | 15350 | 4.2 | 8.09E-36 | 8.09E-36 | 7.45E-36 |
| GOTERM_CC_DIRECT | GO:0005759~mitochondrial matrix | 41 | 6.8 | 1.73E-21 | 572 | 232 | 21777 | 6.7 | 8.01E-19 | 4.01E-19 | 3.88E-19 |
| UP_KW_PTM | KW-0007~Acetylation | 111 | 18.4 | 1.08E-09 | 276 | 2005 | 8410 | 1.7 | 1.84E-08 | 1.95E-08 | 1.84E-08 |
| ***Annotation Cluster 2*** | ***Enrichment Score: 6.78*** | | | | | | | | | | |
| KEGG_PATHWAY | rno01200:Carbon metabolism | 20 | 3.3 | 1.51E-08 | 301 | 133 | 9971 | 5.0 | 4.19E-06 | 1.40E-06 | 1.24E-06 |
| KEGG_PATHWAY | rno00020:Citrate cycle (TCA cycle) | 10 | 1.7 | 2.73E-07 | 301 | 32 | 9971 | 10.4 | 7.60E-05 | 1.52E-05 | 1.35E-05 |
| GOTERM_BP_DIRECT | GO:0006099~tricarboxylic acid cycle | 10 | 1.7 | 3.58E-07 | 561 | 36 | 20691 | 10.2 | 9.79E-04 | 1.96E-04 | 1.95E-04 |
| UP_KW_BIOLOGICAL_PROCESS | KW-0816~Tricarboxylic acid cycle | 9 | 1.5 | 5.33E-07 | 260 | 25 | 8336 | 11.5 | 4.96E-05 | 4.96E-05 | 4.69E-05 |
| ***Annotation Cluster 3*** | ***Enrichment Score: 6.59*** | | | | | | | | | | |
| GOTERM_BP_DIRECT | GO:0006635~fatty acid beta-oxidation | 14 | 2.3 | 8.80E-10 | 561 | 52 | 20691 | 9.9 | 2.40E-06 | 2.40E-06 | 2.39E-06 |
| KEGG_PATHWAY | rno00071:Fatty acid degradation | 12 | 2.0 | 2.00E-07 | 301 | 50 | 9971 | 8.0 | 5.56E-05 | 1.39E-05 | 1.23E-05 |
| UP_KW_BIOLOGICAL_PROCESS | KW-0276~Fatty acid metabolism | 18 | 3.0 | 1.72E-06 | 260 | 142 | 8336 | 4.1 | 1.60E-04 | 7.99E-05 | 7.56E-05 |
| KEGG_PATHWAY | rno01212:Fatty acid metabolism | 11 | 1.8 | 1.41E-05 | 301 | 62 | 9971 | 5.9 | 0.00390157 | 3.26E-04 | 2.89E-04 |

**Table S2. GO ontology enrichment analysis for upregulated genes in the diaphragm of *Gaa*$^{-/-}$ rats.** The top three clusters are presented. Each row represents an enriched GO term or pathway, including details on the number and percentage of genes from the list associated with each term, enrichment significance (p-value), fold enrichment, and the statistical corrections applied (Bonferroni, Benjamini, and false discovery rate or FDR). % is the percentage of genes that are annotated to a specific GO term; "List.Total" stands for the total number of DE genes; "Pop.Hits" stands for the number of background population genes associated with the listed ID of gene ontology term; "Pop.Total" stands for the total number of background population genes. “Fold Enrichment” represents the ratio of genes from that belong to the specific GO term, compared to the expected proportion of genes annotated to that term in the whole genome.

| Category | Term | Count | % | P value | List Total | Pop Hits | Pop Total | Fold Enrichment | Bonferroni | Benjamini | FDR |
|---|---|---|---|---|---|---|---|---|---|---|---|
| *Annotation Cluster 1* | *Enrichment Score: 4.49* | | | | | | | | | | |
| UP_SEQ_FEATURE | MOTIF:Prevents secretion from ER | 10 | 3.9 | 4.96E-10 | 251 | 40 | 22265 | 22.2 | 3.06E-07 | 3.06E-07 | 3.01E-07 |
| GOTERM_CC_DIRECT | GO:0034663~endoplasmic reticulum chaperone complex | 7 | 2.7 | 3.14E-09 | 247 | 13 | 21777 | 47.5 | 1.18E-06 | 5.89E-07 | 5.18E-07 |
| GOTERM_CC_DIRECT | GO:0005788~endoplasmic reticulum lumen | 11 | 4.3 | 4.20E-08 | 247 | 85 | 21777 | 11.4 | 1.57E-05 | 5.25E-06 | 4.62E-06 |
| GOTERM_CC_DIRECT | GO:0042470~melanosome | 9 | 3.5 | 1.14E-05 | 247 | 94 | 21777 | 8.4 | 0.004269665 | 4.75E-04 | 4.18E-04 |
| GOTERM_CC_DIRECT | GO:0005790~smooth endoplasmic reticulum | 6 | 2.3 | 1.68E-05 | 247 | 29 | 21777 | 18.2 | 0.006282115 | 6.30E-04 | 5.55E-04 |
| UP_KW_MOLECULAR_FUNCTION | KW-0143~Chaperone | 12 | 4.7 | 1.77E-05 | 143 | 174 | 10805 | 5.2 | 0.001149627 | 5.75E-04 | 5.75E-04 |
| GOTERM_BP_DIRECT | GO:0006457~protein folding | 10 | 3.9 | 4.15E-05 | 240 | 142 | 20691 | 6.1 | 0.080039717 | 0.010598756 | 0.010487968 |
| KEGG_PATHWAY | rno04141:Protein processing in endoplasmic reticulum | 11 | 4.3 | 7.16E-04 | 161 | 183 | 9971 | 3.7 | 0.162239607 | 0.014746644 | 0.013194366 |
| GOTERM_BP_DIRECT | GO:0034976~response to endoplasmic reticulum stress | 6 | 2.3 | 0.010834833 | 240 | 115 | 20691 | 4.5 | 1 | 0.403036317 | 0.398823394 |
| GOTERM_MF_DIRECT | GO:0051082~unfolded protein binding | 6 | 2.3 | 0.011380098 | 234 | 111 | 19220 | 4.4 | 0.997833637 | 0.225916013 | 0.219172251 |
| GOTERM_MF_DIRECT | GO:0140662~ATP-dependent protein folding chaperone | 4 | 1.6 | 0.017168728 | 234 | 45 | 19220 | 7.3 | 0.999906948 | 0.278312429 | 0.270004596 |
| *Annotation Cluster 2* | *Enrichment Score: 3.05* | | | | | | | | | | |
| GOTERM_CC_DIRECT | GO:0030027~lamellipodium | 11 | 4.3 | 9.67E-05 | 247 | 200 | 21777 | 4.8 | 0.035629484 | 0.002591282 | 0.002280328 |
| KEGG_PATHWAY | rno04530:Tight junction | 11 | 4.3 | 4.02E-04 | 161 | 170 | 9971 | 4.0 | 0.094465325 | 0.009920978 | 0.008876664 |
| KEGG_PATHWAY | rno04670:Leukocyte transendothelial migration | 8 | 3.1 | 0.003140718 | 161 | 120 | 9971 | 4.1 | 0.54020664 | 0.048484833 | 0.043381167 |
| KEGG_PATHWAY | rno04810:Regulation of actin cytoskeleton | 11 | 4.3 | 0.005117202 | 161 | 239 | 9971 | 2.9 | 0.718379264 | 0.066523632 | 0.059521144 |
| *Annotation Cluster 3* | *Enrichment Score: 2.99* | | | | | | | | | | |
| GOTERM_BP_DIRECT | GO:0001916~positive regulation of T cell mediated cytotoxicity | 10 | 3.9 | 2.57E-07 | 240 | 77 | 20691 | 11.2 | 5.16E-04 | 5.16E-04 | 5.10E-04 |
| KEGG_PATHWAY | rno04612:Antigen processing and presentation | 11 | 4.3 | 1.18E-06 | 161 | 87 | 9971 | 7.8 | 2.92E-04 | 1.46E-04 | 1.31E-04 |
| KEGG_PATHWAY | rno05416:Viral myocarditis | 10 | 3.9 | 2.41E-05 | 161 | 97 | 9971 | 6.4 | 0.005934661 | 0.001488067 | 0.001331429 |
| INTERPRO | IPR011161:MHC_I-like_Ag-recog | 7 | 2.7 | 2.96E-05 | 254 | 55 | 23169 | 11.6 | 0.025073298 | 0.025392613 | 0.025007876 |
| GOTERM_CC_DIRECT | GO:0042612~MHC class I protein complex | 6 | 2.3 | 3.24E-05 | 247 | 33 | 21777 | 16.0 | 0.012065711 | 0.001103536 | 9.71E-04 |
| KEGG_PATHWAY | rno05332:Graft-versus-host disease | 8 | 3.1 | 4.07E-05 | 161 | 59 | 9971 | 8.4 | 0.00999676 | 0.002009372 | 0.001797859 |
| UP_KW_CELLULAR_COMPONENT | KW-0490~MHC I | 6 | 2.3 | 5.45E-05 | 195 | 33 | 15350 | 14.3 | 0.001851839 | 9.27E-04 | 6.54E-04 |
| UP_KW_BIOLOGICAL_PROCESS | KW-0391~Immunity | 19 | 7.4 | 5.75E-05 | 101 | 536 | 8336 | 2.9 | 0.003733295 | 0.003740173 | 0.003682632 |
| INTERPRO | IPR003597:Ig_C1-set | 8 | 3.1 | 9.49E-05 | 254 | 97 | 23169 | 7.5 | 0.078206825 | 0.040715269 | 0.040098371 |
| GOTERM_MF_DIRECT | GO:0005102~signaling receptor binding | 16 | 6.2 | 1.17E-04 | 234 | 400 | 19220 | 3.3 | 0.060606909 | 0.010419602 | 0.010108569 |
| KEGG_PATHWAY | rno04514:Cell adhesion molecules | 12 | 4.7 | 1.33E-04 | 161 | 178 | 9971 | 4.2 | 0.032234304 | 0.004095387 | 0.003664294 |
| INTERPRO | IPR037055:MHC_I-like_Ag-recog_sf | 7 | 2.7 | 1.48E-04 | 254 | 73 | 23169 | 8.7 | 0.118915072 | 0.042197306 | 0.041557953 |
| GOTERM_CC_DIRECT | GO:0030670~phagocytic vesicle membrane | 7 | 2.7 | 1.91E-04 | 247 | 74 | 21777 | 8.3 | 0.068999505 | 0.003971592 | 0.003495001 |
| INTERPRO | IPR001039:MHC_I_a_a1/a2 | 6 | 2.3 | 2.33E-04 | 254 | 51 | 23169 | 10.7 | 0.180994868 | 0.049910425 | 0.049154206 |
| SMART | SM00407:IGc1 | 8 | 3.1 | 3.20E-04 | 150 | 97 | 11099 | 6.1 | 0.046826771 | 0.047950954 | 0.047631281 |
| INTERPRO | IPR011162:MHC_I/II-like_Ag-recog | 7 | 2.7 | 3.62E-04 | 254 | 86 | 23169 | 7.4 | 0.267142492 | 0.050251238 | 0.049489855 |
| KEGG_PATHWAY | rno05169:Epstein-Barr virus infection | 13 | 5.1 | 3.64E-04 | 161 | 233 | 9971 | 3.5 | 0.08607987 | 0.009920978 | 0.008876664 |
| INTERPRO | IPR003006:Ig/MHC_CS | 7 | 2.7 | 4.10E-04 | 254 | 88 | 23169 | 7.3 | 0.296600867 | 0.050251238 | 0.049489855 |
| GOTERM_BP_DIRECT | GO:0002476~antigen processing and presentation of endogenous pe | 6 | 2.3 | 5.94E-04 | 240 | 59 | 20691 | 8.8 | 0.69712 | 0.069745187 | 0.069016143 |
| GOTERM_BP_DIRECT | GO:0002486~antigen processing and presentation of endogenous pe | 6 | 2.3 | 5.94E-04 | 240 | 59 | 20691 | 8.8 | 0.69712 | 0.069745187 | 0.069016143 |
| KEGG_PATHWAY | rno05163:Human cytomegalovirus infection | 13 | 5.1 | 8.73E-04 | 161 | 257 | 9971 | 3.1 | 0.193959321 | 0.016578998 | 0.014833841 |
| KEGG_PATHWAY | rno05167:Kaposi sarcoma-associated herpesvirus infection | 12 | 4.7 | 9.85E-04 | 161 | 225 | 9971 | 3.3 | 0.216134923 | 0.017385598 | 0.015555535 |
| INTERPRO | IPR010579:MHC_I_a_C | 4 | 1.6 | 0.001278447 | 254 | 20 | 23169 | 18.2 | 0.666332192 | 0.099718873 | 0.098207981 |
| GOTERM_BP_DIRECT | GO:0045953~negative regulation of natural killer cell mediated cytot | 4 | 1.6 | 0.001286183 | 240 | 19 | 20691 | 18.2 | 0.924746373 | 0.112345262 | 0.111170921 |
| GOTERM_CC_DIRECT | GO:0098553~lumenal side of endoplasmic reticulum membrane | 5 | 1.9 | 0.001387698 | 247 | 43 | 21777 | 10.3 | 0.405924135 | 0.018585244 | 0.016355015 |
| GOTERM_MF_DIRECT | GO:0042605~peptide antigen binding | 7 | 2.7 | 0.001942897 | 234 | 107 | 19220 | 5.4 | 0.647394638 | 0.074385202 | 0.072164748 |
| KEGG_PATHWAY | rno05330:Allograft rejection | 6 | 2.3 | 0.002680856 | 161 | 60 | 9971 | 6.2 | 0.484728092 | 0.044144757 | 0.039497941 |
| KEGG_PATHWAY | rno04940:Type I diabetes mellitus | 6 | 2.3 | 0.004618912 | 161 | 68 | 9971 | 5.5 | 0.681302901 | 0.063381735 | 0.056709973 |
| KEGG_PATHWAY | rno05320:Autoimmune thyroid disease | 6 | 2.3 | 0.006611561 | 161 | 74 | 9971 | 5.0 | 0.805724501 | 0.071002417 | 0.063528479 |
| KEGG_PATHWAY | rno04145:Phagosome | 9 | 3.5 | 0.012490152 | 161 | 193 | 9971 | 2.9 | 0.955152877 | 0.117596286 | 0.10521773 |
| GOTERM_BP_DIRECT | GO:0006955~immune response | 12 | 4.7 | 0.014121423 | 240 | 442 | 20691 | 2.3 | 1 | 0.457009803 | 0.452232697 |

**Table S3. GO ontology enrichment analysis for downregulated genes in the heart of *Gaa*$^{-/-}$ rats.** The top three clusters are presented. Each row represents an enriched GO term or pathway, including details on the number and percentage of genes from the list associated with each term, enrichment significance (p-value), fold enrichment, and the statistical corrections applied (Bonferroni, Benjamini, and false discovery rate or FDR). % is the percentage of genes that are annotated to a specific GO term; "List.Total" stands for the total number of DE genes; "Pop.Hits" stands for the number of background population genes associated with the listed ID of gene ontology term; "Pop.Total" stands for the total number of background population genes. “Fold Enrichment” represents the ratio of genes from that belong to the specific GO term, compared to the expected proportion of genes annotated to that term in the whole genome.

| Category | Term | Count | % | P value | List Total | Pop Hits | Pop Total | Fold Enrichment | Bonferroni | Benjamini | FDR |
|---|---|---|---|---|---|---|---|---|---|---|---|
| ***Annotation Cluster 1*** | ***Enrichment Score: 2.3403422777677676*** | | | | | | | | | | |
| UP_SEQ_FEATURE | TOPO_DOM:Cytoplasmic | 18 | 21.7 | 8.67903E-05 | 74 | 1865 | 22265 | 2.9 | 0.024348209 | 0.024648458 | 0.024474878 |
| UP_KW_BIOLOGICAL_PROCESS | KW-0406~Ion transport | 10 | 12.0 | 0.00018152 | 31 | 602 | 8336 | 4.5 | 0.005792314 | 0.005808627 | 0.005264069 |
| UP_KW_LIGAND | KW-0630~Potassium | 5 | 6.0 | 0.001371627 | 23 | 139 | 6101 | 9.5 | 0.017685138 | 0.017831148 | 0.017831148 |
| GOTERM_BP_DIRECT | GO:0071805~potassium ion transmembrane transport | 5 | 6.0 | 0.002266721 | 75 | 154 | 20691 | 9.0 | 0.809643015 | 1 | 1 |
| UP_SEQ_FEATURE | TOPO_DOM:Extracellular | 12 | 14.5 | 0.005258105 | 74 | 1385 | 22265 | 2.6 | 0.776253524 | 0.276576799 | 0.274629075 |
| UP_KW_BIOLOGICAL_PROCESS | KW-0633~Potassium transport | 4 | 4.8 | 0.008108659 | 31 | 116 | 8336 | 9.3 | 0.229360686 | 0.129738552 | 0.117575562 |
| UP_KW_BIOLOGICAL_PROCESS | KW-0813~Transport | 12 | 14.5 | 0.04233378 | 31 | 1798 | 8336 | 1.8 | 0.749473659 | 0.338670241 | 0.306919906 |
| ***Annotation Cluster 2*** | ***Enrichment Score: 1.9685792755872202*** | | | | | | | | | | |
| KEGG_PATHWAY | rno05207:Chemical carcinogenesis - receptor activation | 7 | 8.4 | 0.00033038 | 42 | 231 | 9971 | 7.2 | 0.045522619 | 0.046583637 | 0.046583637 |
| KEGG_PATHWAY | rno05206:MicroRNAs in cancer | 6 | 7.2 | 0.006863091 | 42 | 295 | 9971 | 4.8 | 0.621307873 | 0.423399436 | 0.423399436 |
| GOTERM_BP_DIRECT | GO:0060291~long-term synaptic potentiation | 4 | 4.8 | 0.021703128 | 75 | 166 | 20691 | 6.6 | 0.999999892 | 1 | 1 |
| GOTERM_CC_DIRECT | GO:0016442~RISC complex | 3 | 3.6 | 0.037937834 | 77 | 88 | 21777 | 9.6 | 0.996604834 | 1 | 1 |
| ***Annotation Cluster 3*** | ***Enrichment Score: 1.8033339369549384*** | | | | | | | | | | |
| UP_SEQ_FEATURE | TOPO_DOM:Cytoplasmic | 18 | 21.7 | 8.67903E-05 | 74 | 1865 | 22265 | 2.9 | 0.024348209 | 0.024648458 | 0.024474878 |
| UP_KW_DOMAIN | KW-1133~Transmembrane helix | 29 | 34.9 | 0.008025748 | 54 | 4782 | 13371 | 1.5 | 0.120964548 | 0.065258818 | 0.065258818 |
| UP_KW_DOMAIN | KW-0812~Transmembrane | 32 | 38.6 | 0.008157352 | 54 | 5509 | 13371 | 1.4 | 0.122828619 | 0.065258818 | 0.065258818 |
| UP_SEQ_FEATURE | TRANSMEM:Helical | 31 | 37.3 | 0.025163064 | 74 | 6621 | 22265 | 1.4 | 0.999281079 | 0.595525858 | 0.591332014 |

**Table S4. GO ontology enrichment analysis for upregulated genes in the heart of *Gaa*$^{-/-}$ rats.** The top three clusters are presented. Each row represents an enriched GO term or pathway, including details on the number and percentage of genes from the list associated with each term, enrichment significance (p-value), fold enrichment, and the statistical corrections applied (Bonferroni, Benjamini, and false discovery rate or FDR). % is the percentage of genes that are annotated to a specific GO term; "List.Total" stands for the total number of DE genes; "Pop.Hits" stands for the number of background population genes associated with the listed ID of gene ontology term; "Pop.Total" stands for the total number of background population genes. “Fold Enrichment” represents the ratio of genes from that belong to the specific GO term, compared to the expected proportion of genes annotated to that term in the whole genome.

| Category | Term | Count | % | P value | List Total | Pop Hits | Pop Total | Fold Enrichment | Bonferroni | Benjamini | FDR |
|---|---|---|---|---|---|---|---|---|---|---|---|
| ***Annotation Cluster 1*** | ***Enrichment Score: 3.26*** | | | | | | | | | | |
| UP_KW_BIOLOGICAL_PROCESS | KW-0391~Immunity | 12 | 9.6 | 0.000377725 | 53 | 536 | 8336 | 3.5 | 0.018712546 | 0.018886274 | 0.018508548 |
| GOTERM_BP_DIRECT | GO:0045087~innate immune response | 10 | 8 | 0.000465494 | 117 | 405 | 20691 | 4.4 | 0.386688841 | 0.079186934 | 0.078055692 |
| UP_KW_BIOLOGICAL_PROCESS | KW-0399~Innate immunity | 8 | 6.4 | 0.00093025 | 53 | 254 | 8336 | 5.0 | 0.045468049 | 0.023256257 | 0.022791132 |
| ***Annotation Cluster 2*** | ***Enrichment Score: 2.97*** | | | | | | | | | | |
| GOTERM_CC_DIRECT | GO:0005615~extracellular space | 25 | 20 | 5.42909E-05 | 115 | 1936 | 21777 | 2.4 | 0.011497658 | 0.011563953 | 0.011129626 |
| UP_KW_DOMAIN | KW-0732~Signal | 38 | 30.4 | 0.002172846 | 75 | 4458 | 13371 | 1.5 | 0.023643327 | 0.023901311 | 0.023901311 |
| UP_KW_PTM | KW-1015~Disulfide bond | 30 | 24 | 0.01073154 | 71 | 2359 | 8410 | 1.5 | 0.167583323 | 0.193167719 | 0.193167719 |
| ***Annotation Cluster 3*** | ***Enrichment Score: 2.48*** | | | | | | | | | | |
| GOTERM_BP_DIRECT | GO:0045071~negative regulation of viral genome replication | 5 | 4 | 0.000163554 | 117 | 49 | 20691 | 18.0 | 0.157806372 | 0.057243761 | 0.056425993 |
| GOTERM_BP_DIRECT | GO:0009615~response to virus | 5 | 4 | 0.002431424 | 117 | 100 | 20691 | 8.8 | 0.922393521 | 0.189941018 | 0.187227575 |
| GOTERM_BP_DIRECT | GO:0035455~response to interferon-alpha | 3 | 2.4 | 0.003552066 | 117 | 16 | 20691 | 33.2 | 0.97615807 | 0.196298408 | 0.193494145 |
| KEGG_PATHWAY | rno03250:Viral life cycle - HIV-1 | 4 | 3.2 | 0.009778328 | 70 | 64 | 9971 | 8.9 | 0.843890728 | 0.264014862 | 0.257030342 |
| UP_KW_BIOLOGICAL_PROCESS | KW-0051~Antiviral defense | 3 | 2.4 | 0.029284449 | 53 | 43 | 8336 | 11.0 | 0.773744394 | 0.366055617 | 0.358734505 |
